# Prior learning shapes effort choice, revealing valence asymmetries and disruption in anhedonia

**DOI:** 10.1101/2025.07.24.666522

**Authors:** Calum Guinea, Bowen Xiao, Rebecca P. Lawson

## Abstract

Goal directed behaviour requires the integration of learned beliefs about the world with decisions about whether an outcome is worth the effort required to obtain it. Although value learning and effort-based choice are central to motivated behaviour, they are typically examined in isolation. Here we introduce a novel behavioural paradigm that directly links probabilistic associative learning with effortful decision making, for the first time allowing trial by trial assessment of how learned beliefs guide willingness to exert effort.

In a large sample of participants (n=252), individuals reliably acquired stimulus-outcome associations but showed substantial variability in learning fidelity. Effort allocation was not determined by objective reward probabilities, but was better predicted by participants subjective beliefs, demonstrating that effortful action depends on internal value representations rather than external contingencies alone. Reward and loss contexts showed asymmetric effects on effort motivation, revealing a valence specific dissociation in belief driven action.

Critically, higher levels of anhedonia were associated with reduced willingness to engage in effortful behaviour and a weaker coupling between learned value and action, indicating a disruption in the integration of belief and motivation. Together, these findings identify a mechanistic link between learning and effort expenditure and suggest that motivational impairments in psychiatric symptoms may arise from failures to translate learned value into action.

## Introduction

The way we allocate effort in the present is inherently shaped by what we have learned about the value of available options in the past. Yet, despite this intrinsic connection, research on learning and effort-based decision-making has largely evolved in parallel, forming substantial but predominantly separate bodies of literature in cognitive neuroscience and related fields. To date, no existing work has systematically examined how prior learning influences subsequent effort-based decision-making.

Decision-making can be understood as a form of cost-benefit analysis, where people weigh potential rewards against associated costs. Principal among these costs is effort: be it physical or cognitive, each decision we make is likely to incur some effort cost. For example, visiting a new café might require walking two miles. While the absolute effort cost (i.e., the distance) is fixed, individuals differ in their sensitivity to that cost (i.e., how aversive they find the walk) and in how much they value the potential reward (i.e., how enjoyable they expect the café to be). Simply put, the decision to visit the café depends on whether the anticipated reward justifies the required effort.

Early studies established that humans generally seek to minimise effort whenever possible^1,2^. This aversion applies to both physical^3^ and cognitive effort^4^, but can be overridden when effort leads to reward. Notably, in some cognitive tasks, effort itself can be intrinsically rewarding or preferred over rest^5^. When access to rewards is effort-dependent, we conceptualise the reward’s subjective value as being discounted based on the effort required to obtain it^6^. Importantly, this effort discounting occurs even when rewards are hypothetical, such as virtual apples^7^ or virtual money^8^. While effort discounting is observed in the general population, it is steeper in individuals with psychiatric symptoms such as apathy and anhedonia^9,10^.

Willingness to exert effort for reward is most commonly assessed using effort-based decision-making tasks^10–14^. Physical effort has been measured experimentally in several ways, including repeated pressing of keys on a keyboard^10,15^, but most commonly, using grip force, where the effort requirements in the task are defined as a proportion of a person’s maximum grip strength (Maximum Voluntary Contraction, MVC)^11–14^. Typically, these tasks then ask participants to choose whether to exert different levels of effort for different amounts of reward such that effort exertion is deterministically related to reward (see ^15,16^ for exceptions). However, in naturalistic decision making, exerting effort is often probabilistic and does not guarantee the receipt of reward (i.e. just because we exert effort to visit the new café, it does not guarantee that our experience of the café will be positive). Our tasks must therefore account for the *likelihood* of getting the desired outcome or avoiding the undesired one. To reflect the probabilistic relationships between effort and outcomes, some experimental designs choose to cue the probability of receiving reward at the start of each trial^15–17^. While this enhances experimental control, it comes at the cost of overlooking a crucial individual difference – namely, that people may vary in the subjective beliefs they acquire about the uncertain relationship between effort and its outcomes. These differences in subjective beliefs can arise from differences in learning, which produce different experiences of uncertainty. Extending our example, you may feel uncertain about a new café, as you have little prior knowledge about the quality of its coffee. Yet uncertainty applies even to familiar choices – your usual café might still disappoint despite past good experiences. How we learn about this uncertainty and use what we have learned to guide our choices is therefore both central to naturalistic effort choice behaviour and psychiatrically relevant^18–22^.

Evident throughout the literature on learning is the idea that people learn differently, even from identical experiences. Computational models highlight variability in learning rates - some people update beliefs quickly, others slowly^18,23,24^. A simpler way to measure people’s learning is through explicit choices or preferences, having people learn probabilistic relationships between stimuli and rewards, then selecting between the stimuli in a “tournament”. This approach provides a detailed measure of how accurately they learned those initial probabilities^25–27^. Such differences matter for effort-based decisions – if someone overestimates reward probability (e.g. believing that a 70% chance of reward is actually 80%), they may be more willing to exert effort. Conversely, someone who underestimates reward probability might be less willing to exert effort. Learning accuracy is therefore important to effort-based choice, yet effort decision-making studies often treat reward as deterministic, such that if the effort requirement on a given trial is met, then reward is obtained^14^. In reality, effort decisions hinge on learning accuracy, reward sensitivity and motivation. Without measuring and independently manipulating these variables, we cannot pinpoint why people differ in effort exertion. This may be particularly important in the context of mental health conditions where diagnoses affecting learning and effort decision-making are highly co-occurring^28^.

Finally, most of the research on effort-based decision-making has tended to focus on effort for reward^3,10,11,13–15,29^, though the vigour of effort to prevent negative outcomes has been studied previously^30^. Our willingness to exert effort to avoid negative outcomes has rarely been studied, despite the fact that avoidance and apathy are characteristic of mood and anxiety disorders^6,31^. Valence-driven differences in learning from prediction errors are well-documented^32^, and loss aversion is a robust feature of economic decision-making - losing £100 feels worse than gaining £100 feels good^33^. Some studies have investigated willingness to exert effort for loss; in one example, participants were required to exert either high or low effort for appetitive or aversive stimuli^34^. Here, effort exertion is deterministically related to the outcomes; this exertion was not calibrated to the individual (as is common in grip force paradigms^14,35^ and participants can never entirely avoid exerting effort. In a study of prosocial effort, the grip force task was used to compare participants’ willingness to exert effort to obtain reward or avoid loss for oneself or for another person^35^. Again, effort exertion was deterministically related to the outcomes participants received and motivation to obtain reward and avoid loss was not compared. As with reward, these losses may be probabilistic and require the decision-maker to track or infer this probability, alterations in the tracking and estimation of uncertainty have regularly been associated with mood disorders^36^. Given the literature’s emphasis on effort for rewards, whether humans avoid effort to prevent losses in the same way they pursue effort for gains remains an open question.

### Psychiatric relevance of learning and effort-based choice

Both learning and effort-based decision-making are of established psychiatric and neuropsychological interest. Anxiety has been associated with elevated or improper adaptation of learning rates^18,37^ and, different types of anxiety drive distinct biases in safety versus threat learning^38^. Misestimating uncertainty can reinforce avoidance – anxious individuals may overgeneralise from negative experiences, exerting excessive effort to avoid potential threats. Core features of anxiety like rumination, are effortful because of the processes they recruit, like mental replay of unpleasant experiences^39^ and persist despite the cost they incur, which suggests that anxiety may be associated with an increased willingness to exert effort. While avoidance characteristically defines anxiety disorders, evidence also links anxiety to increased willingness to exert cognitive effort in reward-based tasks^40^. Recent literature has found that different types of anxiety have different behavioural effects on learning^38^ and effects separable from depression^41^. Given the heterogeneity of experience in mood disorders^42^, its impact on physical effort remains little understood.

Anhedonia and apathy also shape effort decisions^7,10,15,16,40^. Anhedonia, often tied to depression but present across many psychiatric and neurological diagnoses^6^, reduces effort motivation and blunts sensitivity to reward probability and outcome magnitude^15,16,40^. It has also been argued that anhedonia reflects impaired Pavlovian biases (i.e. action to approach reward and suppress action to avoid) rather than altered reward or punishment sensitivity^43^. Fatigue, a symptom that is likewise implicated across psychiatric diagnoses, further complicates the specificity of anhedonia and apathy’s effects on effort decision-making. Fatigue frequently co-occurs with both, and all three can manifest as lack of motivation and reduced energy levels. Computational modelling has shown that fatigue affects people’s ability to persist in effortful tasks, potentially mimicking or amplifying the effects of apathy and anhedonia^14^. Distinguishing these constructs is crucial for understanding how different psychiatric symptoms contribute to effort-based decision-making.

Effort-related impairments likely arise from multiple mechanisms – limiting the utility of diagnostic labels. This reflects a near-ubiquitous issue with current psychiatric nosology, for which several alternatives have been proposed to simplify or improve upon it (e.g. p-factor^44^ and the RDoC matrix^45^). Transdiagnostic factor modelling, which takes a large number of self-reported questionnaire responses about mental health symptoms and identifies common structure amongst them, has been used to both simplify and identify commonalities across psychiatric diagnoses^46^. Using this approach, the anxious-depression factor and the social withdrawal factor are both associated with the avoidance of mental effort^47^. Such dimensional approaches, then, might be fruitful in disentangling the impact of blurred psychiatric constructs on learning and effort decision-making.

Here we propose that across individuals, the fidelity of a person’s learning, their sensitivity to reward and their sensitivity to effort may have convergent effects on effort-based decision-making. To arbitrate between these possible causes, we designed a novel task that combines Pavlovian learning with effort-based decision-making. This allows us to dissociate the contributions of learning, reward sensitivity and effort sensitivity on effort decision-making, offering a clearer picture of how these factors shape decision-making in anxiety, anhedonia and apathy. First, in the learning phase (**Figure 1A**), participants (n=252) learned probabilistic associations between eight different aliens with varying outcome probabilities (0.25, 0.75) and magnitudes of reward/loss (+ 40 points, +160 points, - 40 points, - 160 points), reporting objective estimates of the outcome probability on each trial. We then assessed how well participants could discriminate between pairs of these stimuli in a “tournament” phase (**Figure 1B**). Finally, participants chose to ‘accept’ or ‘reject’ the opportunity to exert effort (keypresses) to approach or avoid one of the eight aliens they had previously learned about (effort decision-making phase; **Figure 1C**). Effort levels (low - 60%, medium - 75%, high - 90%) were adjusted to each participant’s maximum number of presses in three calibration trials at the beginning of the experiment.

**Figure 1:**
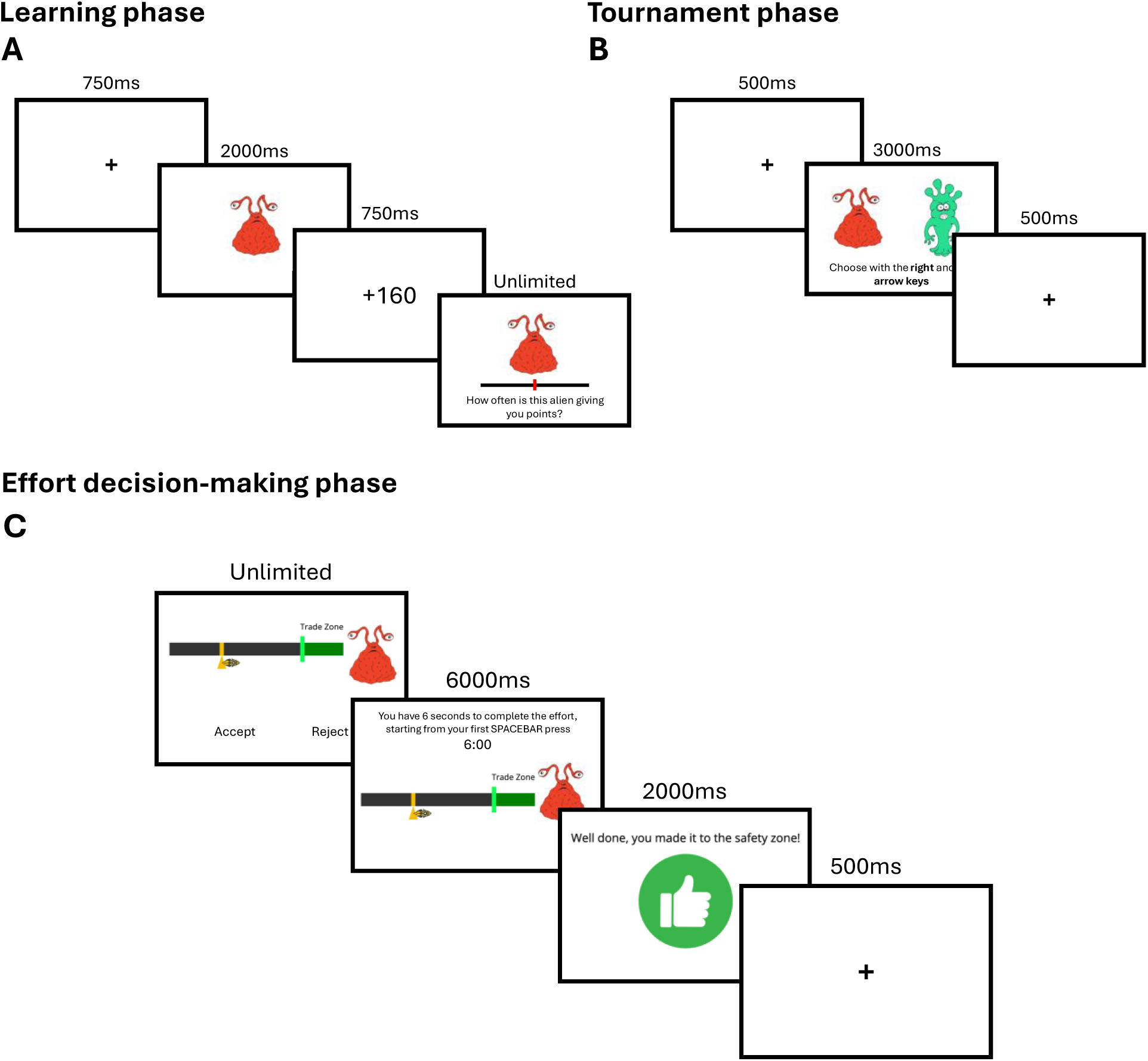
Learning, tournament and effort-decision making phases of the experiment. **(A)** An example trial from the learning phase of the experiment. Participants first view a fixation cross, then one of the eight stimuli is shown on screen before the outcome of the trial appears on the next screen. For reward stimuli, when the alien gives points, a positive tone is played when the amount appears on screen and for loss trials, a negative tone is played when the alien takes points. For both alien types, a neutral tone is played when the outcome is zero. At the end of each trial, participants are asked to estimate the percentage of trials that the alien is giving or taking points from them, this estimate is reported using a slider. **(B)** An example trial from the tournament. Participants see a brief fixation cross and are then presented with two of the eight learned stimuli and asked to make a choice between them in 3000ms using the left and right arrow keys on their keyboard. If no choice is made this is recorded as no response. If participants make a choice in under 3000ms the next pair of stimuli appear on the screen after a brief delay. **(C)** An example trial from the effort decision-making phase where the participant accepts the offer and meets/exceeds the effort requirement. On each effort trial, participants are presented with one of the eight stimuli they learned about previously and an effort amount visualised as the distance from a Trade/Safety zone on a horizontal bar. Participants are required to accept or reject based on the effort required and what they remember about the outcome probability and magnitude associated with the given stimulus. When accepting the offer, participants are taken to a screen which includes a countdown timer that counts down to zero from 6000ms as soon as the participant makes the first spacebar press. Rejecting the effort offer takes participants to a waiting screen that also last for 6000ms.

## Results

### Effort calibration

All participants, except one, comfortably exceeded the required minimum of 19 presses in one of their three calibration trials. The average number of keypresses across the sample was 45, with a standard deviation of 11.004 and a good range of individual differences captured, min = 19, max = 78 (**Figure S1**).

### Measures of learning

Several robust behavioural indicators suggest that participants were, on average, able to learn the outcome probabilities of the eight different aliens.

#### Trial-by-trial estimates of stimulus outcome probability

In the learning phase of the experiment, participants’ trial-by-trial estimates of each alien’s outcome probability trended towards the true outcome probability (**Figure 2A**). However, there was significant variation across the sample. There were substantial individual differences in learning accuracy, with some participants overestimating the true outcome probability and others tending to underestimate the true outcome probabilities (**Figure 2B and 2C**).

**Figure 2:**
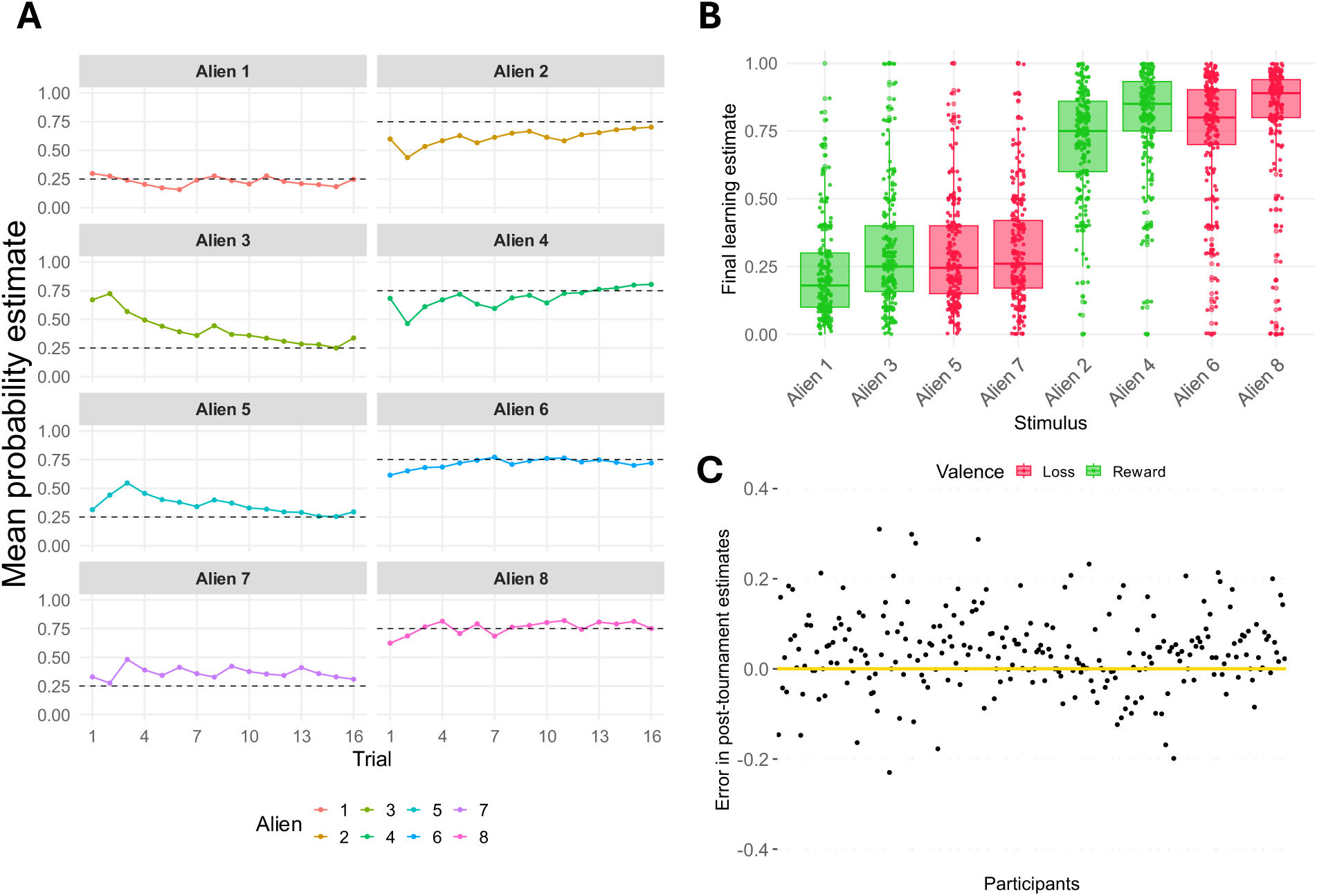
Summary of performance in the learning phase of the task. **(A)** The average of all participants’ probability estimates across the learning trials. Here we take the mean rating of all participants’ estimates for each stimulus on each trial (coloured lines) to visualise the trajectory of participants’ learning and how, across trials, their ratings tend toward the true outcome probability (dashed black line). **(B)** Box and scatter plots of participants’ final learning estimate for each stimulus. The stimuli have been ordered to group the low probability (p = 0.25) and high probability (p = 0.75) stimuli. Green indicates reward stimuli and red indicates loss stimuli. Participants tended towards the objective outcome probability but there was also notable variability in participants’ estimates. **(C)** Here we take the mean post-tournament probability estimate of each participant and subtract the true average outcome probability from it (0.5), such that participants who on average, overestimate the true outcome probabilities are above the yellow line. This average error indicates there is significant variability in learning fidelity across participants.

#### Post-task outcome probability ratings

At the end of the learning, tournament and effort-choice tasks, participants gave ‘final’ explicit ratings of the outcome probability for all eight aliens. This provides a measure of what was learned about the stimuli and whether it changed across time (**Figure S2**). After initial learning, participants’ average estimates of the outcome probabilities did not significantly change across time points (F (2, 502) = 0.851), Greenhouse-Geisser corrected p = 0.428; see **Figure S2**), suggesting that learning remained stable. Comparing participants’ average estimates against the true average probabilities of the stimuli indicated that, at the group level, they generally correctly approximated the true outcome probabilities (**Figure 2C; Table S1**).

#### Tournament performance as a measure of learning

After the learning phase, participants completed a binary-choice tournament task which enabled a more detailed assessment of participants’ preferences amongst the learned stimuli. Overall participants made accurate choices (mean = 78%, SD = 10.95) and were good at approaching the best alien when it was present in a choice pair (mean = 85%, SD = 17.57) as well as avoiding the worst stimulus when it was present in a pair (mean = 82%, SD = 18.89). A Bayesian t-test provided support for the null hypothesis that there was no difference in participants’ approach and avoid accuracy (BF_10_ = 0.304). When choices were more difficult, with pairs containing two stimuli of the same valence, participants were less accurate for both reward (mean = 65%, SD = 20.19) and loss (mean = 62%, SD = 17.84). Again, a Bayesian t-test indicated there was weak evidence for the alternative hypothesis that reward and loss accuracy were different (BF_10_ = 0.19). See **Figure S3**.

### Effort decision-making

In the effort decision making task, participants accepted a high percentage of effort offers overall (mean = 75%, SD = 18.81) and tended to accept higher effort offers (mean = 59%, SD = 28.82) less than medium effort offers (mean = 77%, SD = 22.41) and medium effort offers less than low effort offers (mean = 91%, SD = 14.55). Overall participants’ effort choices were sensitive to effort level (F (1, 251) = 353.6, p < 0.001), outcome magnitude (F (1, 251) = 71.82, p < 0.001), outcome probability (F (1, 251) = 207.3, p < 0.001) and valence (F (1, 251) = 15.81, p < 0.001).

There was a significant effort by outcome probability interaction (F (2, 502) = 75.79, p < 0.001), indicating that participants accepted more effort for higher probability outcomes than for lower probability outcomes. A significant outcome magnitude by outcome probability interaction (F (1, 251) = 16.91, p < 0.001), suggesting participants accepted more offers for higher probability and higher outcome magnitude stimuli. There was a significant outcome magnitude by valence interaction (F (1, 251) = 29.4, p < 0.001), showing that participants accepted similar amounts of effort for low magnitude reward and loss stimuli but accepted more effort for larger magnitude rewards than losses.

Lastly, there was a significant four-way interaction between effort, outcome probability, outcome magnitude and valence (F (1, 502) = 13.56, p < 0.001). Across all effort levels, participants accepted more effort for reward than for loss. Generally, they discriminated more precisely between high and low probability as well as high and low magnitude reward stimuli than loss stimuli. For example, participants accepted effort for a similar amount of low and high magnitude stimuli on low effort trials, regardless of outcome probability. Overall, these results suggest that the levels of effort, probability, outcome magnitude and valence were well calibrated in this part of the experiment (see **Figure 3** for 3D bar plots representing how the probability of accepting changes across these variables), with main effects and interactions as expected based on prior literature (Treadway et al., 2009; Treadway et al., 2012).

**Figure 3:**
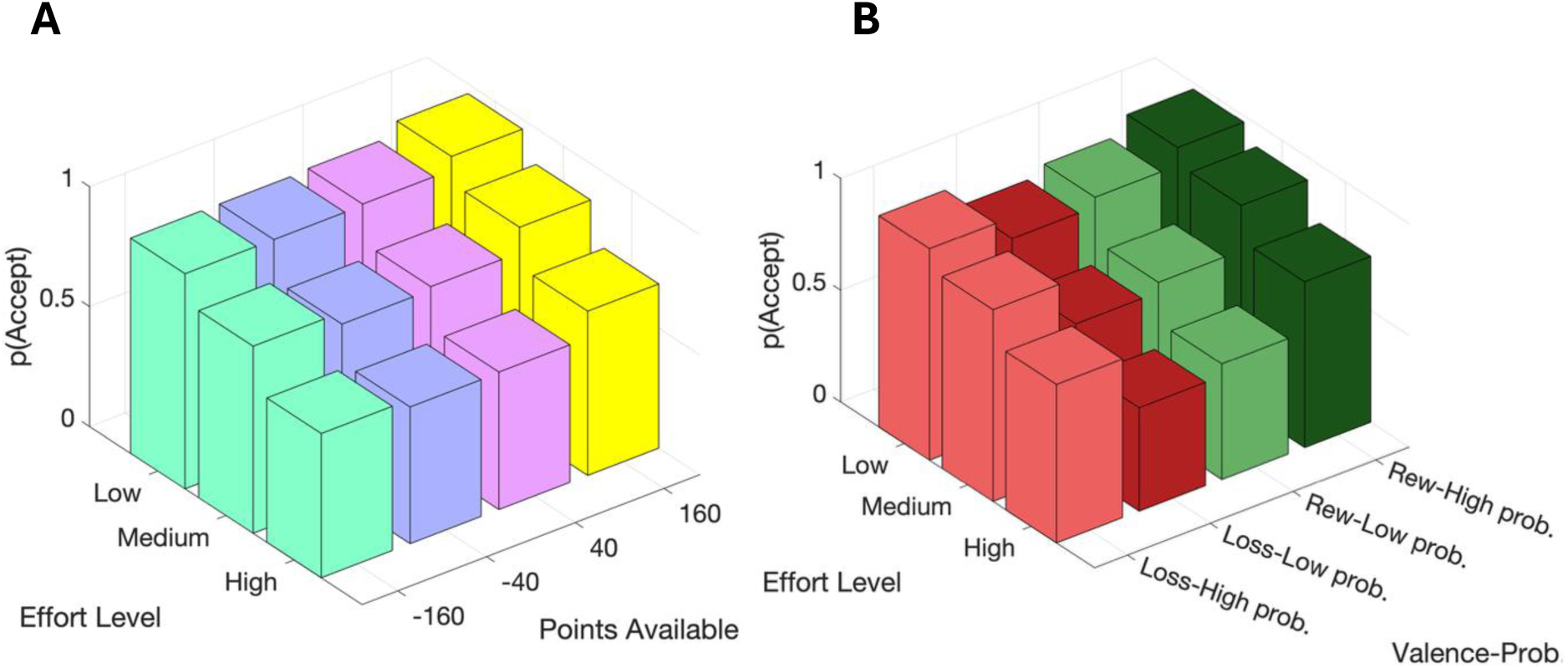
Probability of accepting as a function of effort level, outcome magnitude and valence-probability. **(A)** A 3D bar plot showing how the probability of accepting effort varies as a function of effort level and the amount of points associated with a stimulus. The probability of accepting was highest for the highest outcome magnitude stimuli combined with the lowest effort and lowest for the low magnitude loss stimuli combined with the highest effort level. For each level of outcome magnitude, p(accept) decreased as effort level increased. **(B)** A 3D bar plot showing how the probability of accepting effort varies as a function of effort level and outcome probability for both reward and loss aliens. The probability of accepting was highest for the high probability reward stimuli combined with the lowest effort level and lowest for the low magnitude loss stimuli combined with the highest effort level. For each stimulus valence and outcome probability combination, p(accept) decreased as effort level increased.

### Effort choices are sensitive to prior learning

After establishing the key behavioural features of the learning, tournament, and effort-based decision-making phases of our experiment, we now examine our primary hypothesis: that individual differences in learning the outcome probabilities of each stimulus influence participants’ subsequent effort choices. First, we fit a logistic regression (M0) that assumes all participants learn equally and perfectly about the probability of the aliens they encountered. Full details of all predictors in the model are outlined in the methods section below (see also **Table S2)**. Overall, the model was a significant improvement over the null (X^2^ = 4221.8, p < 0.001, McFadden R^2^ = 0.156). As expected, effort level (β = −7.004, p < 0.001) and objective outcome probability (β = 2.58, p < 0.001) were significant predictors of effort choice but in opposite directions; with increasing effort level reducing the odds of accepting an effort offer and increasing outcome probability increasing the odds of accepting. Likewise, outcome magnitude was a significant predictor of effort choice (β = 0.774, p < 0.001); the odds of accepting increased with increased outcome magnitude. We also observed small but significant effects of learning order (β = −0.102, p = 0.002), where learning about the loss aliens first was associated with decreased odds of accepting an effort offer, and an effect of valence (β = 0.224, p < 0.001), where reward aliens were associated with increased odds of accepting. Of the demographic variables, only gender was a significant predictor (p < 0.001), with small but significant differences largely driven by non-binary participants and those not reporting a gender identity (see **Table S2**).

As M0 assumes participants learn accurately and identically to one another, we next compared a suite of models with different individualised measures of participants’ learning as predictors of effort-choice. Full details of all models considered can be found in **Supplementary Table 3**. Crucially, all models containing individual signatures of learning significantly improved the prediction of participants’ effort-choice over and above M0 (see **Supplementary Table 3** for a comparison of models containing individualised learning measures and **Supplementary Figure 4** for a correlation matrix of the relationships amongst these learning measures), but the winning model substituted participants’ *post-tournament probability estimates* in place of the objective outcome probabilities.

Post-tournament probability estimates were a significant predictor (β = 6.01, SE = 0.37, Wald stat. < 0.001) of effort choice, for every unit of increase in participants’ probability estimates, the log odds of accepting an effort offer increased by 6.01. Outcome magnitude was a small, positive and significantly non-zero predictor of effort choice, with the log odds of accepting an effort offer increasing by 0.34 (SE = 0.16, Wald stat. p = 0.035) for every unit of increase in outcome magnitude. Stimulus valence was also a positive and significantly non-zero predictor of effort choice, with the log odds of accepting an effort offer increasing by 0.39 (SE = 0.09, Wald stat. p < 0.001) for reward stimuli. By contrast, effort level (β = −12.78, SE = 0.65, Wald stat. p < 0.001) was a negative and significantly non-zero predictor, for every unit of increase in effort level the log odds of accepting an effort offer decreased by −12.78. See **Figure 4** for a visualisation of the fixed effects and slope estimates from the best-fitting model M1.

**Figure 4:**
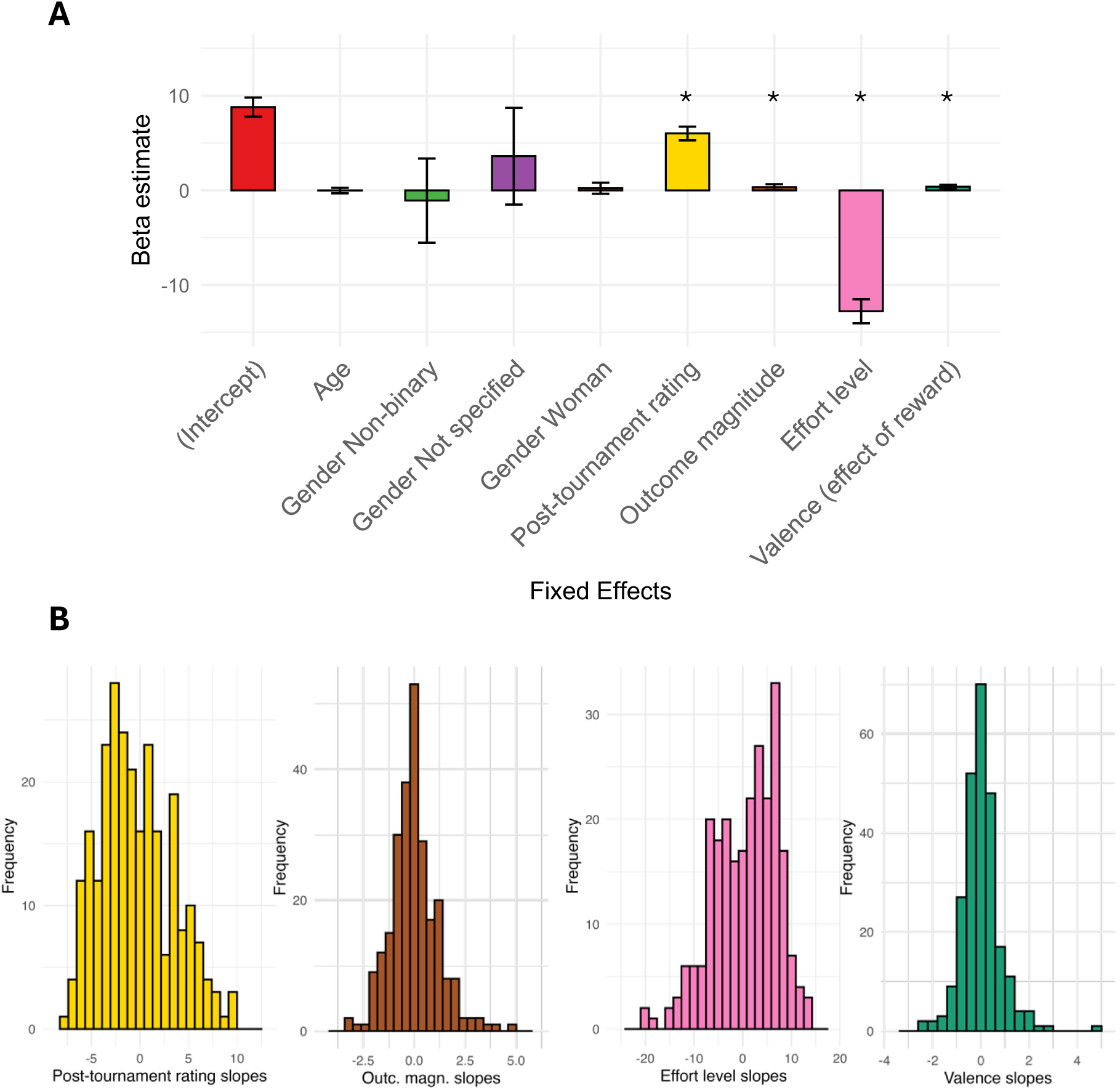
Bar plot of fixed effect estimates and histograms of the random effects from M1. **(A)** Bar plot of the fixed effect estimates from Model 1 (M1), the winning model amongst those fit to compare the performance of individualised learning measures against the objective outcome probabilities. Each bar represents the fixed effect estimate for a predictor from the winning model (M1). Error bars represent the 95% confidence interval estimates for the predictor. Unlike in Model 0 (M0), in the GLMMs, none of the demographic predictors were significantly non-zero predictors of trial-by-trial effort choice. Post-tournament rating, outcome magnitude, effort level and stimulus valence were all significantly non-zero predictors (p < 0.05). **(B)** Histograms of the random slope estimates for each of the predictors in the winning model, Model 1 (M1). For example, where the fixed effect estimate for effort level is zero, each participant’s slope estimate for the effect of effort level on choice is expressed as a difference from zero. Therefore, participants with a positive slope estimate for effort level are more sensitive to effort level when making effort choices. The slopes for each random effect are approximately normally distributed.

### The effect of self-report mental health measures on effort choice

Over and above demonstrating that a model accounting for the individual signatures of learning improves the prediction of participants’ choices, we sought to understand whether transdiagnostic and self-report measures of mental health symptoms also predicted participants’ effort choices. **Figure S5** shows the distribution of scores on the SHAPS, FAS and the transdiagnostic factor dimensions across the sample. Accordingly, we fit three additional binomial generalised linear mixed models (GLMMs) by adding the transdiagnostic and self-report mental health measures as fixed effects to the best-fitting model (M1). Model 2 (M2) added the three transdiagnostic symptom measures, Model 3 (M3) added the sum score of the SHAPS and Model 4 (M4) added the FAS. AIC scores suggest that Model 3 (M3), which included the SHAPS sum score, was the best performing of these models (**Table S4**) and crucially, also outperformed Model 1 (M1). Participants post-tournament probability estimates (β = 5.98, SE = 0.37, Wald stat. p < 0.001), stimulus outcome magnitude (β = 0.34, SE = 0.16, Wald stat. p = 0.032), effort level (β = −12.8, SE = 0.65, Wald stat. p < 0.001) and valence (β = 0.4, SE = 0.1, Wald stat. p < 0.001) remain significant in this model. In addition to the significant predictors of the previously reported model, participants’ sum scores on the SHAPS were a significantly non-zero predictor of their effort choices (β = −0.34, SE = 0.15, Wald stat. p = 0.026). The log odds of accepting decreased by 0.34 for every unit of increase in SHAPS sum score (**Figure 5**). In comparison, none of the three transdiagnostic factors or the FAS were significant fixed effects predictors of effort choice in their respective models (see **Table S4** for a comparison of these models containing mental health measures against Model 0 and Model 1). As we identified an effect of the SHAPS on trial-by-trial effort choice, we ran partial correlations controlling for the effects of age and gender. SHAPS scores were significantly negatively correlated with the percentage of medium (τ = −0.086, p = 0.043) and high effort offers accepted (τ = −0.083, p = 0.049). Kendall’s tau was used here because the percentage of effort offers accepted is not normally distributed, and there were a number of participants with the same percentage of offers accepted (e.g. 100% of low effort offers accepted).

**Figure 5:**
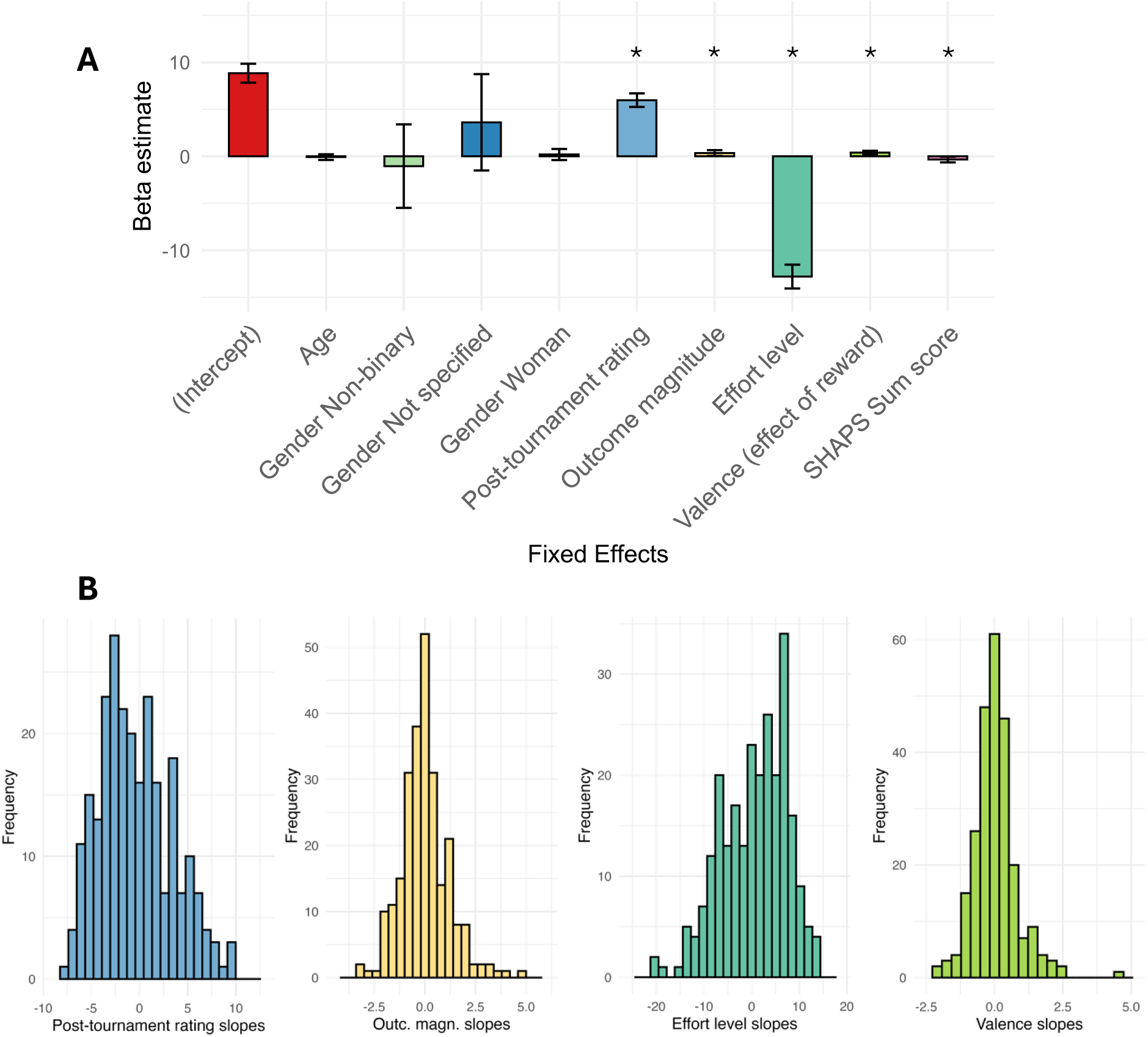
Bar plot of the fixed effects and histograms of the random effects from M3. **(A)** Bar plot of the fixed effect estimates from the overall best-fitting model, which included participants’ SHAPS sum scores as a fixed effect predictor (Model 3). Each bar represents the fixed effect estimate for a predictor from the winning GLME model (Model 3), error bars are 95% confidence intervals. All predictors are significant (p < 0.05). **(B)** Histograms of the random slopes estimated for each participant from the winning model (Model 4). The slopes fit for each participant are expressed as a difference from zero, where zero is the fixed effect estimate for the predictor. So, for post-tournament rating, a participant with a positive slope estimate suggests that the participant’s effort choices are more strongly influenced by their own probability estimate of the stimulus. For each predictor, the slope estimates are approximately normally distributed.

### The interaction of anhedonia and individualised measures of learning

As our analysis of participants’ trial-by-trial effort choices showed that participants’ scores on the SHAPS were a significant fixed effect predictor of their effort choices, we conducted some additional exploratory analyses to assess whether SHAPS scores significantly interact with any of the decision variables included in our winning model (M3; see **Supplementary Materials** for full details of these exploratory models). Only the interaction between participants’ post-tournament probability estimates and SHAPS scores was significant (β = −0.71, SE = 0.33, p = 0.0303; **Supplementary Materials EM1**). This suggests that the effort choices of participants with higher anhedonia are less sensitive to their own estimates of the outcome probabilities. Specifically, compared to participants reporting no or low anhedonia, those with high anhedonia are less willing to accept effort as their estimates of the stimulus outcome probability increase (**Figure 6**; see also **Figure S6** for a visualisation of how the probability of accepting varies with SHAPS scores across effort levels).

**Figure 6:**
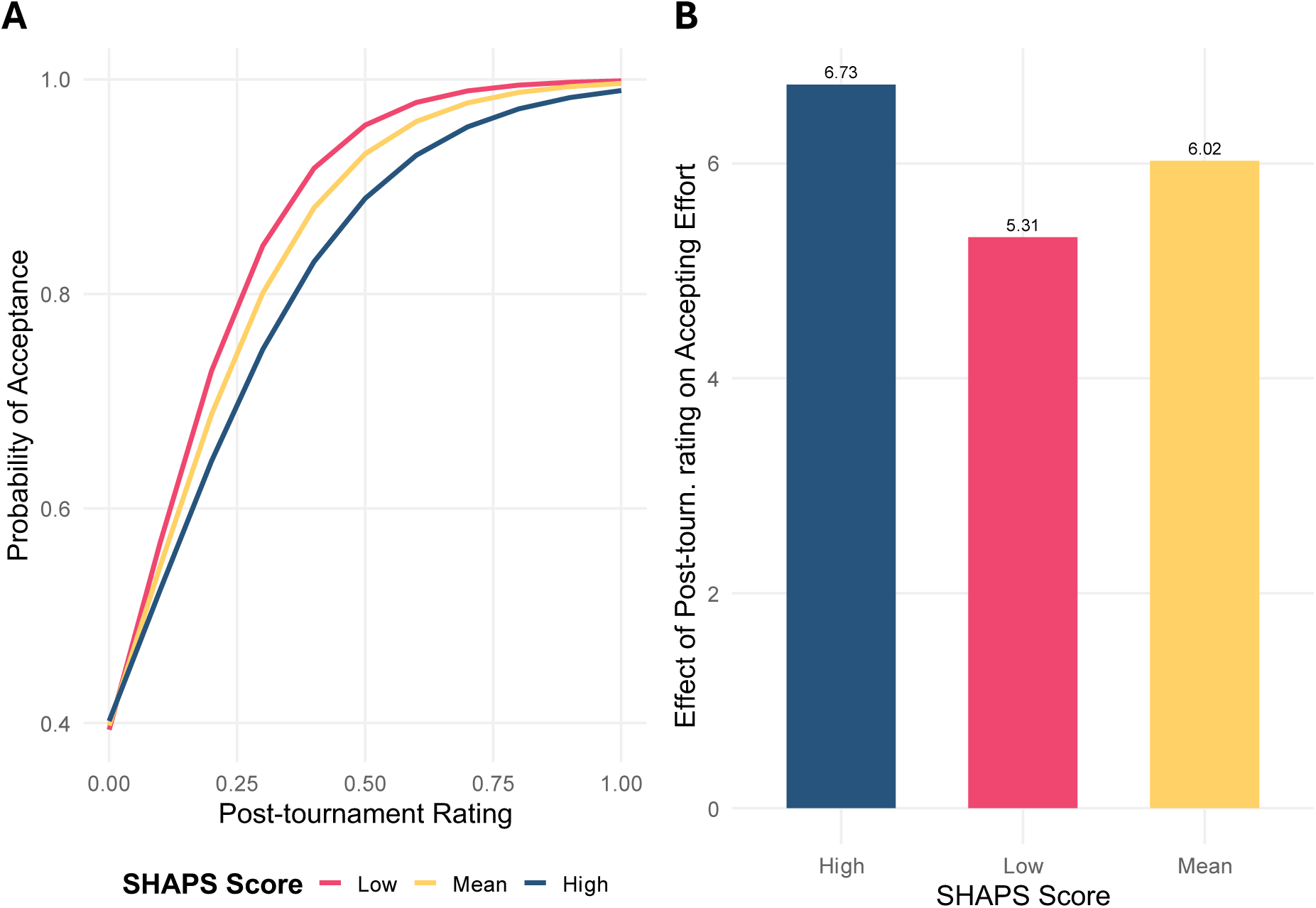
The interaction bet een SHAPS scores and participants’ post-tournament probability estimates. **(A)** Predictions from our winning interaction model (EM1) that visualise the effect of the interaction between participants’ post-tournament probability estimates and SHAPS scores on the probability of accepting effort offers. Compared to participants scoring lower on the SHAPS (cream colour), high SHAPS scorers’ (red colour) effort choices are more weakly related to their estimates of the stimuli’s outcome probability. Lower scorers on the SHAPS show a sharper increase in probability of accepting as their estimate of the outcome probability increases than participants who score higher on the SHAPS. **(B)** A bar plot of the interaction effect estimates for high, low and mean scorers on the SHAPS. These estimates were calculated by adding and subtracting the estimate for the interaction effect from the fixed effect estimate for participants’ post-tournament estimates from the winning interaction model (EM1).

When participants estimate that the outcome probability of a stimulus is high, all participants are equally likely to accept effort offers. However, as participants’ estimates of the outcome probability decrease, participants scoring high on the SHAPS are less likely to accept effort offers. All other predictors in the model remained significant except the fixed effect of SHAPS scores (β= 0.01, SE = 0.22, p > 0.05)

## Discussion

Learning and effort-decision making are both central to motivated behaviour, yet they are typically studied in isolation. By integrating these processes into a single paradigm, we capture a more ecologically valid model of goal-directed behaviour. Our novel task combined probabilistic associative learning with effort decision-making to understand how these two behaviours interact. Analysis of learning and tournament behaviour clearly demonstrates that participants were able to learn the outcome associations of the stimuli and then use that learning to accurately discriminate amongst them. However, we also observe significant variability in learning across participants. Crucially, trial-by-trial effort decisions were best predicted by participants’ *own* estimates of stimulus-outcome probabilities, rather than the objective probabilities. Moreover, individuals with higher anhedonia scores were less inclined to exert effort and exhibited a weaker alignment between their learned beliefs and their decisions—suggesting a disruption in the translation of expected value into motivated action.

A striking finding was the asymmetry in participants’ willingness to exert effort for potential reward versus avoiding equivalent losses. On average, participants were more inclined to work for a possible gain than to prevent a comparable loss, despite equal expected value. This asymmetry was robust and did not vary with anhedonia, suggesting it reflects a fundamental motivational bias rather than individual differences in hedonic capacity. To date, few studies have investigated how individuals make effort choices to avoid negative outcomes, in these studies successful completion of the effort always leads to reward^34,35^. Here we combine features from existing paradigms, namely: the calibration of effort to the individual and providing participants with the opportunity to avoid effort entirely and extend this work by requiring participants to learn the probabilistic outcomes of effort. Our design combines two major advantages that more closely mirror real-world effort decision-making, first allowing participants to opt out of effort so individuals can choose rest over exertion. Second, by making effort probabilistically related to outcomes, individuals must consider that there is uncertainty about whether their effort exertion will have the expected outcome. These features enhance sensitivity to symptoms like anhedonia, as their expression in naturalistic behaviour may be to opt out of effortful behaviour altogether, yet this opting out could result from reduced motivation, reduced outcome sensitivity or inaccurate learning about this uncertainty. While prior research has overwhelmingly focused on effort in service of obtaining reward, our study broadens the concept of goal-directed behaviour to include the avoidance of loss. Crucially, we find that reward and loss stimuli with equal absolute expected value do not motivate effort exertion equally, here participants were more willing to exert effort to obtain reward than to avoid loss.

Like apathy, anhedonia has been associated with reduced willingness to exert effort for reward^15,16,48^. Consistent with this, we observed that participants who scored higher on self-reported anhedonia were less likely to accept effortful offers on a trial-by-trial basis. Importantly, exploratory analyses revealed a significant interaction between anhedonia and participants’ own estimates of stimulus outcome probability, such that higher reported anhedonia was associated with weaker coupling between their choices and their probability estimates. In other words, compared to those with lower anhedonia, their likelihood of accepting a given effort offer increased less steeply as a function of their perceived probability of an outcome. Accordingly, we report weak but significant negative correlations between self-reported anhedonia and the percentage of both medium and high effort offers. These findings suggest that anhedonia may disrupt the ability to integrate expected value and outcome probability when deciding whether to exert effort, potentially leading to less adaptive motivational behaviour.

To our knowledge, this is the first study to examine the relationship between transdiagnostic mental health factors^49^ and physical effort choice. These factors putatively capture a range of symptoms that are likely related to choice (e.g. items from the Obsessive-Compulsive Inventory)^50^ and effort (e.g. items from the Apathy Evaluation Scale; AES)^51^. Despite this, no previous work has assessed whether such transdiagnostic constructs relate to physical effort choice. One prior study previously reported associations between the anxious-depression and social-withdrawal factors and avoidance of mental effort^47^. Though the anxious-depression factor comprises symptoms related to apathy (Apathy Evaluation Scale)^51^, social anxiety. (Liebowitz Social Anxiety Scale)^52^, depression (Self-rating Depression Scale)^53^ and state and trait anxiety (State-Trait Anxiety Inventory)^54^, it does not necessarily capture anhedonia and here we did not find evidence of similar associations between anxious-depression and *physical* effort.

One possibility is that the heterogeneous symptoms subsumed by these factors exert opposing influences on physical effort-based decision-making, leading to overall null effects. Indeed, recent work has taken a more targeted approach, using factor analysis across a battery of questionnaires specifically designed to deeply phenotype anhedonia and apathy^55^ - symptoms with clearer theoretical links to motivational impairments. Moreover, the impact of symptoms on effort decision-making may be both symptom-specific and domain-specific. Supporting this, recent work assessing participants’ sensitivity to both physical and mental effort costs found that apathy and anhedonia have a domain-specific effect on physical effort sensitivity, while anxiety specifically influenced sensitivity to cognitive effort^40^. These findings underscore the importance of examining specific symptom dimensions, rather than broad transdiagnostic factors, and of distinguishing between different types of effort when investigating motivational impairments in mental health.

Cognitive and physical effort are distinct forms of exertion and may have distinct relationships to psychiatric symptoms^40^. Anxiety has been linked to reduced sensitivity to cognitive effort, potentially due to the high costs associated with symptoms like worry and rumination, which involve excessive mental replay^39^. In contrast, depression-related symptoms are typically studied in relation to physical effort^15^, where they are associated with increased effort discounting^10^. While the cognitive and physical domains may involve distinct mechanisms, they may converge through their accumulation of fatigue – a potential shared cost signal. Recent computational work has further distinguished between two types of fatigue: one that is recoverable through rest, and one that is not^14^. These latent components of fatigue have been shown to closely track participants’ subjective estimates of effort exertion and tiredness^56^. Fatigue thus appears central to understanding effort decision-making. However, in our study we were unable to apply these modelling approaches to estimate individual fatigue parameters, as they rely on patterns in trial-by-trial decision behaviour that were not elicited in our design. Moreover, in our study participants likely experienced and recovered from varying degrees of fatigue across the learning and tournament phases prior to the effort-choice task. While self-reported fatigue was measured, we found no evidence that it influenced effort decisions in this sample, unlike in recent work which has reported similar effects of fatigue and apathy on effort decision-making^9^. Future studies using dynamic fatigue modelling may reveal how individual differences in fatigue shape effort-based decisions across domains.

Though we observed substantial individual variation in learning measures, these differences were not associated with any indicators of mental health. At first this might seem surprising, given that alterations in learning have been reported in association with a range of different psychiatric diagnoses and symptoms^43^. However, the learning task used here was deliberately simple—featuring fixed outcome probabilities, no opportunities for choice, and an emphasis on tracking probabilities rather than magnitudes. This simplicity was important to ensure that participants could reliably acquire the outcome contingencies, as this learning served as a foundation for subsequent effort-based choice. As a result, this learning task was not able to investigate the associations between specific mental health symptoms and learning behaviour seen in the literature, including: the relationship between anhedonia and choice temperature^43^ as participants did not make choices during learning; the improper adaptation of learning rate in anxiety^18^ or internalising psychopathology more generally^57^ as the outcome contingencies were stable for all learned stimuli and, anhedonia’s relationship to reduced reward sensitivity^58^ as the requirement to provide trial-by-trial estimates of stimulus outcome probability likely shifted participants attention away from outcome magnitude. Future studies should test whether such associations might emerge under more complex or uncertain learning conditions, where greater demands are placed on flexibility, valuation, or motivational processes.

By integrating probabilistic associative learning and effort-based decision-making in a single task, our results reveal that participants can learn equally well but differ in how learning informs action, particularly in the context of psychiatric symptoms. This work bridges two previously separate literatures and suggests that anhedonia-specific mechanisms may underlie impairments in goal-directed behaviour.

## Methods

### Participants

In total, 299 participants were recruited using the online recruitment platform Prolific using the following inclusion/exclusion criteria: aged 18-65, fluent English speaker, not colourblind and a Prolific study approval rating above 90%, refer to **Table 1** for a summary of the sample’s demographic information. All these participants provided informed consent at the start of the study. After applying further exclusion criteria, 252 participants were included in the final analyses. 42 participants were excluded for repeatedly failing at least one of the instruction quizzes designed to test participants’ understanding of the learning, tournament and effort decision-making tasks. Five participants were removed because their overall choice accuracy in the tournament task was 50% or lower, indicating random responding. Participants who completed all tasks and questionnaires received £10.50. This study was approved by the Cambridge Psychology Research Ethics Committee (CPREC; Ethics number PRE.2019.110).

**Table 1:**
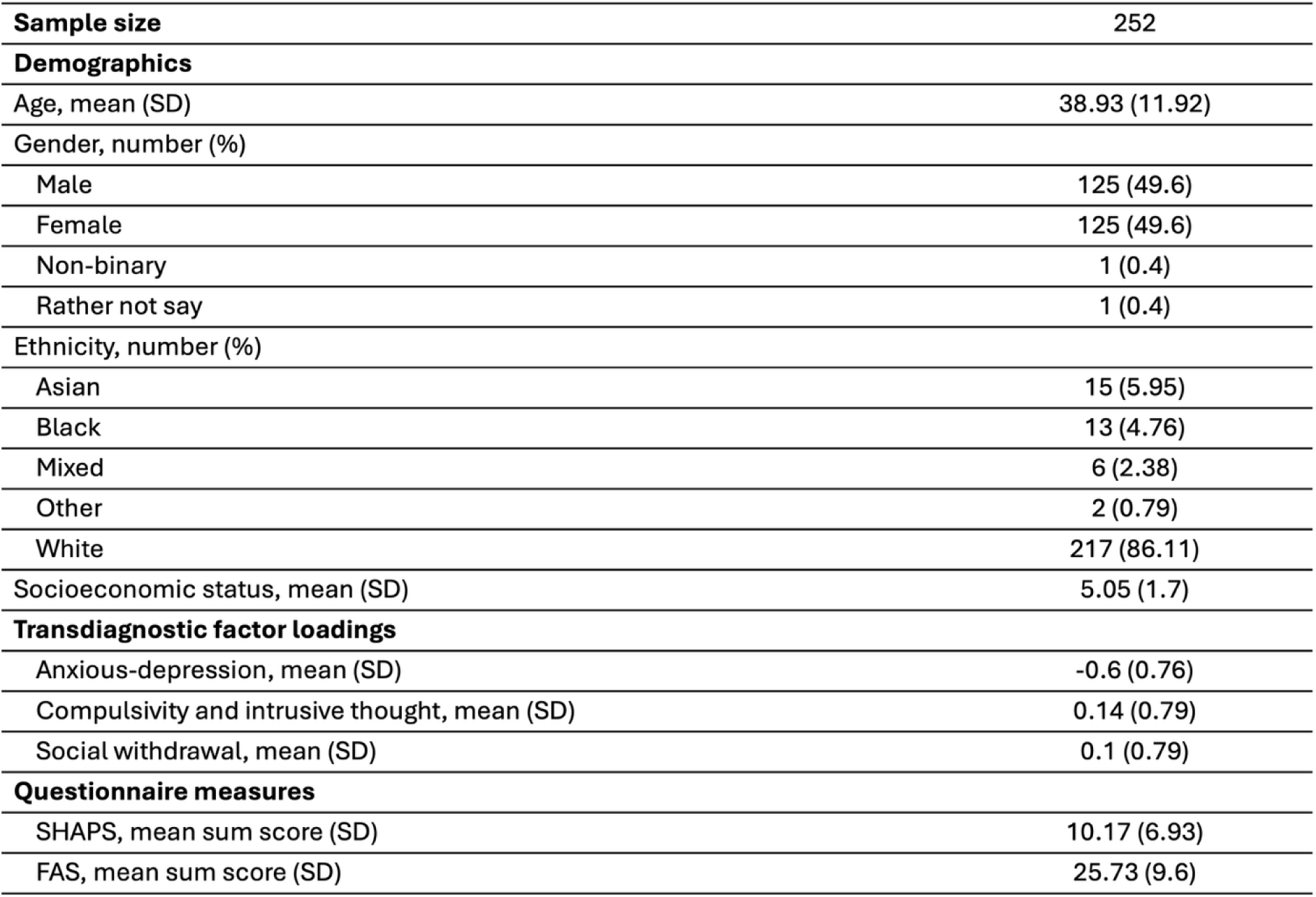
Demographic information of the sample. . Participant loadings onto the three transdiagnostic factors and scores on self-report measures of anhedonia and fatigue.

### Task design

#### Effort calibration

At the start of the experiment, to establish each individual’s maximum capacity for key pressing, participants were asked to press the spacebar on their keyboard using their right and left index fingers as fast as possible in a six second window. With every keypress the participant added a star graphic to the screen and their task was to “fill the sky with stars” to help light the path home for an astronaut. The maximum number of keypresses is a measure comparable to maximum voluntary contraction from grip force tasks^11–14^. This was repeated three times, with their highest number of key presses from the three runs taken forward into the effort decision-making. After each attempt, participants were told how many stars they added to the sky and were encouraged to try and beat it on the next attempt. To prevent participants from minimising their effort, participants need to press at least 19 times to proceed beyond the calibration stage of the experiment.

#### Pavlovian Learning

Participants learned about eight different stimuli (aliens) and their probabilistic association with reward or loss outcomes. Four of the aliens were probabilistically associated with reward and four were probabilistically associated with loss. The outcome probabilities and magnitudes were arranged such that each stimulus was associated with a unique combination of probability and magnitude (see **Supplementary Table 5**). The reward and loss aliens were presented in separate blocks and the trial order in each block was randomised once, such that all participants experienced the same order of trials. Each alien was presented a total of 16 times so participants completed 128 trials across the two blocks. On each trial, participants were shown one of the eight aliens and given an outcome (here points that were not related to participants’ payment for participation in the study). They were then asked to use a slider to estimate the outcome probability that this alien gives you or takes away points (i.e. the reward or loss probability of the alien; see **Figure 1 Panel A** for a schematic of the learning trials). This permitted the explicit measure of trial-by-trial learning across encounters with a given alien, without the need for instrumental responses that might confound subsequent measures of choice in later stages of the experiment. As participants encountered reward and loss aliens in separate blocks, the order of these blocks was counterbalanced across the sample, half of participants learned about the reward aliens first and the other half of participants learned about the loss aliens first.

At the end of the final learning block, participants were asked to provide a final estimate of the reward probability of all eight stimuli.

#### Tournament

After the learning phase, participants were asked to complete a ‘tournament’ in which they made binary decisions between pairs of the eight aliens. All possible pair combinations of the stimuli were presented five times in a randomised order, with all participants experiencing the same trial order. Participants were instructed to choose between the alien pairs based on which one they felt was ‘best’ from the learning phase. Participants had three seconds to respond to each pair using the left and right arrow keys to choose the left and right stimuli respectively. If participants did not respond within the three second limit, this was recorded as no response and the next pair would appear on the screen immediately (see **Figure 1 Panel B**).

At the end of the tournament, participants were again asked to provide a final estimate of the reward probability of all eight stimuli.

#### Effort-based decision-making

Then participants completed an effort-based decision-making task (**Figure 1 Panel C**). On each trial, participants chose to ‘accept’ or ‘reject’ the opportunity to exert effort to approach or avoid one of the eight aliens they had previously learned about. Prior to starting the effort-based decision-making, participants are reminded to press the spacebar with their right and left index fingers as they did during the calibration task. Here effort is defined as the number of keypresses required to move a spaceship to reach a ‘trade’ (approach) or ‘safety’ (avoid) zone (see screen two in **Figure 1 Panel C**). Participants were presented with one of three effort levels (60%, 75% and 90% of each participant’s maximum number of key presses), represented on a horizontal bar in the middle of the screen showing the distance they would be required to travel. For approach trials, the alien was presented next to the ‘trade’ zone, indicating this was an approach trial. Participants were instructed that accepting an effort offer on this trial meant agreeing to exert effort for the chance to trade with the alien located in the ‘trade’ zone. Whereas for avoid trials, the learned stimulus was presented on the left side of the effort bar, indicating that it was an avoid trial. On these trials, participants were instructed that accepting an effort offer here was like taking the opportunity to avoid having points stolen by the alien. Critically, no information was presented about the reward/loss probability or magnitude associated with each alien, requiring participants to make choices based on the effort level and *what they learned about the stimuli previously*.

When participants accepted an effort offer, they progressed to a second screen and had six seconds to meet the effort requirement, the six second countdown started on their first button press. Participants could see their spaceship progress toward the ‘trade’ or ‘safety’ zone and therefore, determine whether they had met the effort requirement based on whether their spaceship had passed the green marker on the effort bar (see screen two in **Figure 1 Panel C**). In both approach and avoid trials, participants were only told whether they had met the effort requirement or not. This prevented participants from continuing to learn about the reward/loss probabilities of the aliens. On trials where participants rejected the effort offer, participants had the opportunity to rest for a duration equivalent to the effort exertion window (six seconds). Participants completed 96 effort choices in total, which ensured that each combination of effort level and stimulus was presented four times in total in a pre-randomised order and the same order for all participants.

At the end of the effort-choice task, participants were again asked to provide a final estimate of the reward probability of all eight stimuli.

#### Transdiagnostic and self-report mental health measures

At the end of the experiment participants completed a shortened version of the transdiagnostic three factor questionnaire^59^ which contains questions relating to apathy, the Snaith-Hamilton Pleasure Scale (SHAPS)^60^, a measure of anhedonia which we calculate as a sum score to increase its fidelity as a predictor^61^ and the Fatigue Assessment Scale (FAS)^62^ to measure fatigue.

### Analysis

#### Measuring learning

With the inclusion of trial-by-trial probability estimates, final post-task probability estimates and tournament performance, there are several ways to quantify how participants learned the outcome probabilities associated with each alien. Analyses were conducted across a range of measures (outlined below) to confirm that, on average, participants were capable of learning the stimulus-outcome probabilities in this task.

##### Mean trial-by-trial probability estimates

here we calculate the mean of a participant’s probability ratings for each alien (16 trials) of the learning task.

##### Final learning probability estimates

here we take the last estimate participants gave for each stimulus during the learning task. The trial order of the learning was randomised for the reward and loss blocks, meaning the final learning trials for each stimulus were not the last eight trials of the learning.

##### Post-task ratings

here we took the explicit rating provided for each stimulus at the end of a given task (e.g. post-learning, post-tournament and post-effort choice). This allowed us to examine whether learning persisted or changed across different phases of the experiment.

##### Tournament performance

To assess choice accuracy, we calculated the expected value (EV) of each alien by multiplying its outcome magnitude (points) by its outcome probability. The correct choice for a given pair was the alien with the higher EV. We also examined approach accuracy (choosing the highest-EV alien when present) and avoid accuracy (avoiding the lowest-EV alien when present)^25^. Additionally, we measured accuracy in choices between only reward or only loss stimuli (‘within-valence’ comparisons).

#### Logistic regression predicting trial-by-trial choice

To first establish that participants’ effort choices were sensitive to the effects of effort, outcome probability and outcome magnitude, a baseline logistic regression was used to predict participants’ trial-by-trial choice data with effort level, outcome probability and outcome magnitude, learning order, stimulus valence, age, gender and socioeconomic status as predictors. Learning order was a binary predictor included to measure a possible effect of block order on effort choice and valence was also a binary predictor included as this design presents a unique opportunity to compare how people make effort choices to approach reward and avoid loss when outcome probabilities and absolute outcome magnitudes are equivalent. Gender uses male (including trans male) as a reference category, the remaining levels were: female (including trans female), non-binary and gender not specified. These models were fit using R (R version 4.4.1; R Core Team, 2024; https://www.R-project.org/) and *stats* package (version 4.4.1; R Core Team, 2024).

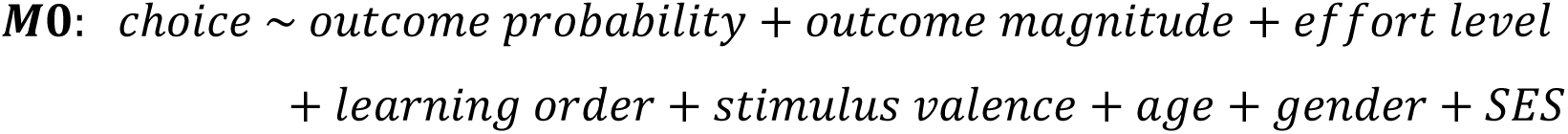

#### Generalised linear mixed effects modelling

The simple logistic regression model above assumes that all participants learn perfectly (i.e. they learn the exact outcome probabilities of each stimulus) and identically to one another (i.e. the logistic regression treats all participants as one participant). To capture individual differences in learning, a set of models which included different individualised measures of participants’ learning were also fit to determine whether these individual measures of learning better explain participants’ effort choices (refer to **Supplementary Table 3** for a full list of models and see the **Supplemental Materials** for details of the measures tested). We allowed formal model comparison to arbitrate between different individual learning signatures as predictors of effort-choice, and report details of the winning model in the results section. Here we used AIC which punishes less for model complexity than BIC to identify winning models but AIC scores, along with BIC scores, log likelihood and deviance are also reported in the **Supplementary Materials**. Bold text indicates where models have changed from the previous model:

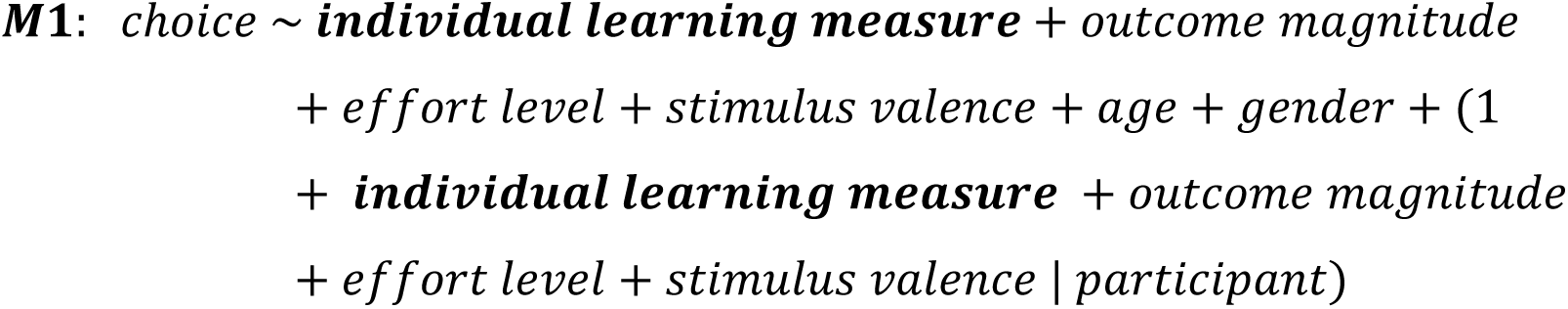

Lastly, in this general population sample, we were interested in whether participants’ effort choices were related to a set of relevant psychiatric dimensions and clinical symptoms (e.g. three transdiagnostic factor scores comprising anxious-depression, compulsivity and intrusive thought and social withdrawal; in addition to anhedonia and fatigue). We therefore fit the following suite of models (bold text indicates where models have changed from the previous model):

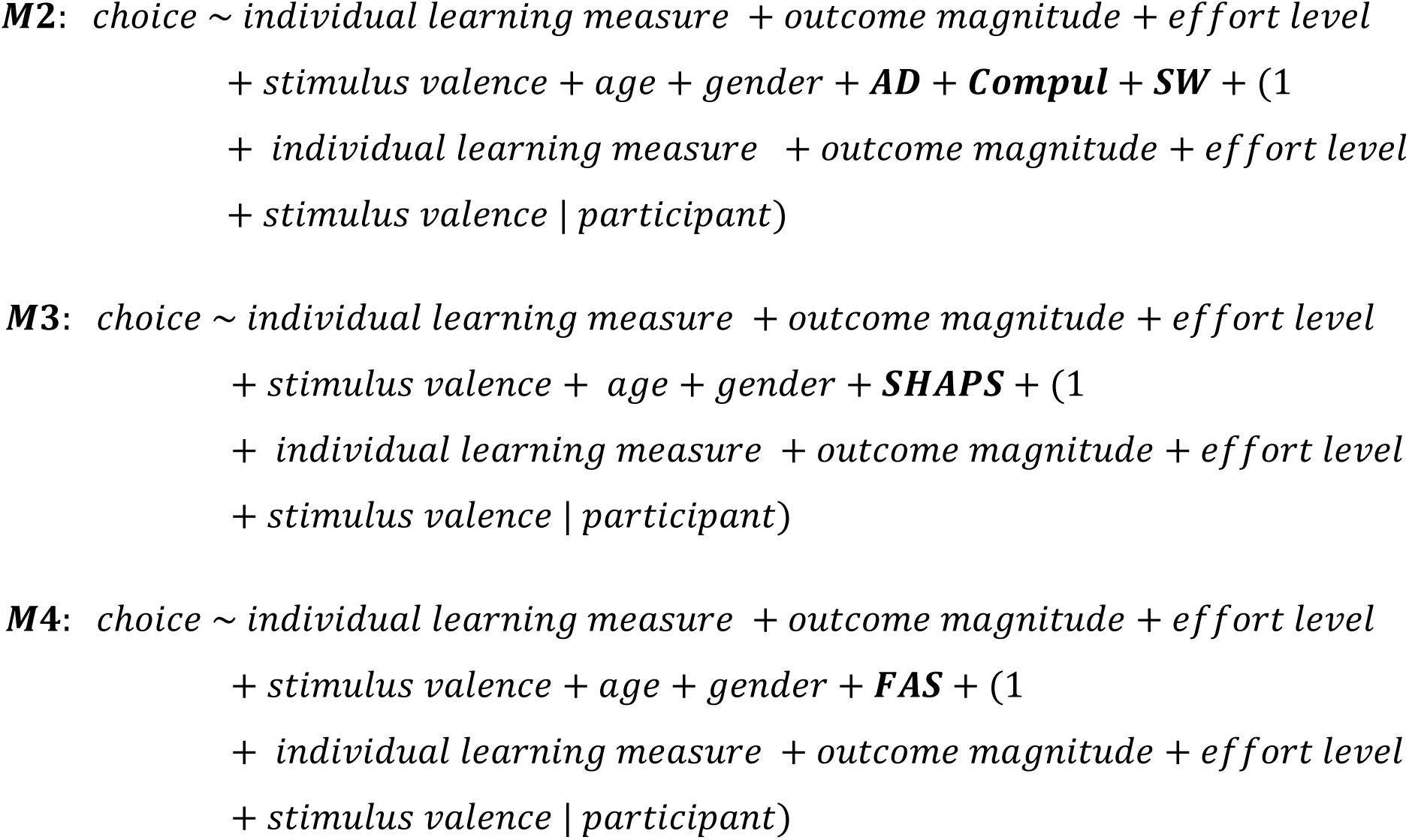

To improve mixed model fitting and to aid interpretation and comparison of estimates generated by these models, questionnaire scores were rescaled and zero-centred. As all values of effort level and outcome probability ranged between 0 and 1, outcome magnitude was min-max normalised with a minimum of 0.2 and a maximum of 0.8, such that the lowest absolute outcome magnitude became 0.2 and the highest absolute outcome magnitude became 0.8 thus preserving the relationship between the original absolute magnitudes of 40 and 160 (i.e. the large outcome magnitude being four times the small outcome magnitude).

## Data availability

The data from this study are available from the corresponding author upon reasonable request and via a public Github repository (https://github.com/Calumlg/lxmAnalysisPub).

## Ethics

This work was approved by the Cambridge Psychology Research Ethics Committee (CPREC), ethics number PRE.2019.110.

## Supplementary Materials

**Supplementary Figure 1:**
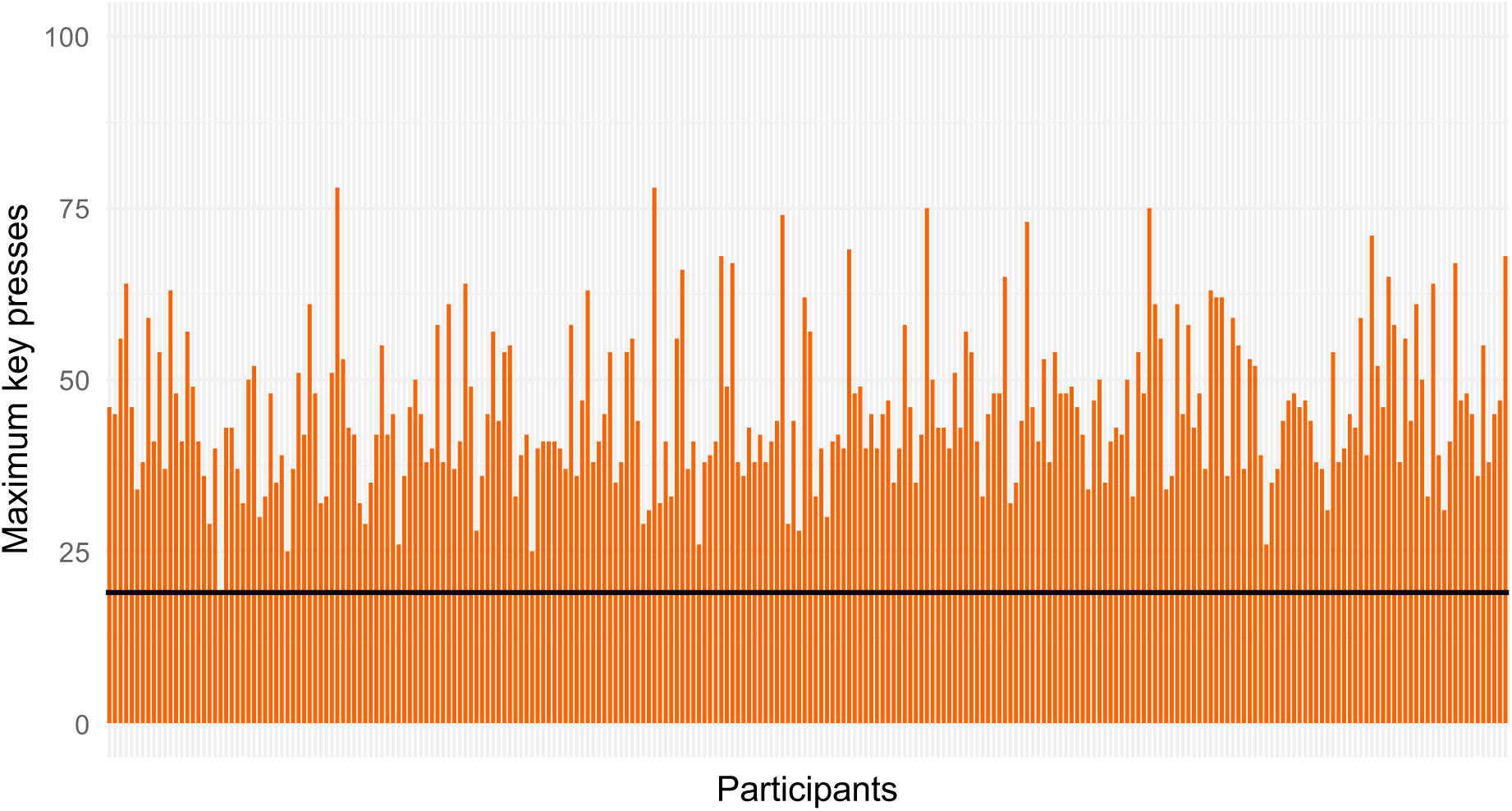
Bar plot of participants’ maximum key presses from the effort calibration. Each bar represents a participant’s maximum number of key presses from the three calibration trials completed at the start of the experiment. The highest number of presses completed in the three calibration trials was taken forward to the effort decision-making task. Only one participant in the final sample had a maximum number of key presses equal to the minimum required to participate in the remainder of the experiment (19 presses).

**Supplementary Figure 2:**
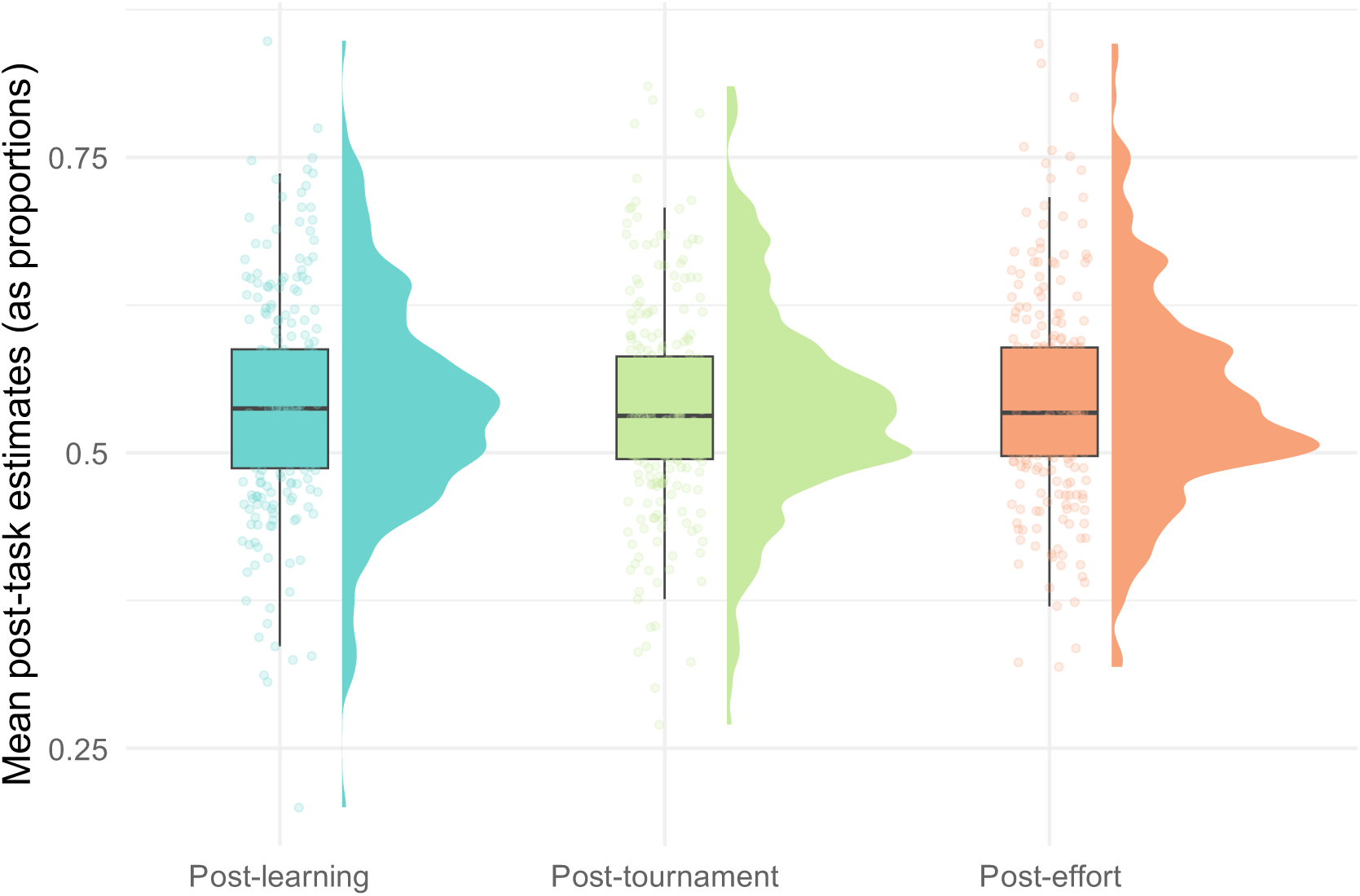
Mean post-task estimates across the experiment. Rain cloud plots represent the distribution of participants’ outcome probability estimates, which remain consistent across the experiment. The boxplot represents the 25th and 75th percentiles, and the middle line represents the median rating. Whiskers are 1.5 times the 25^th^ and 75^th^ percentiles.

**Table S1:**
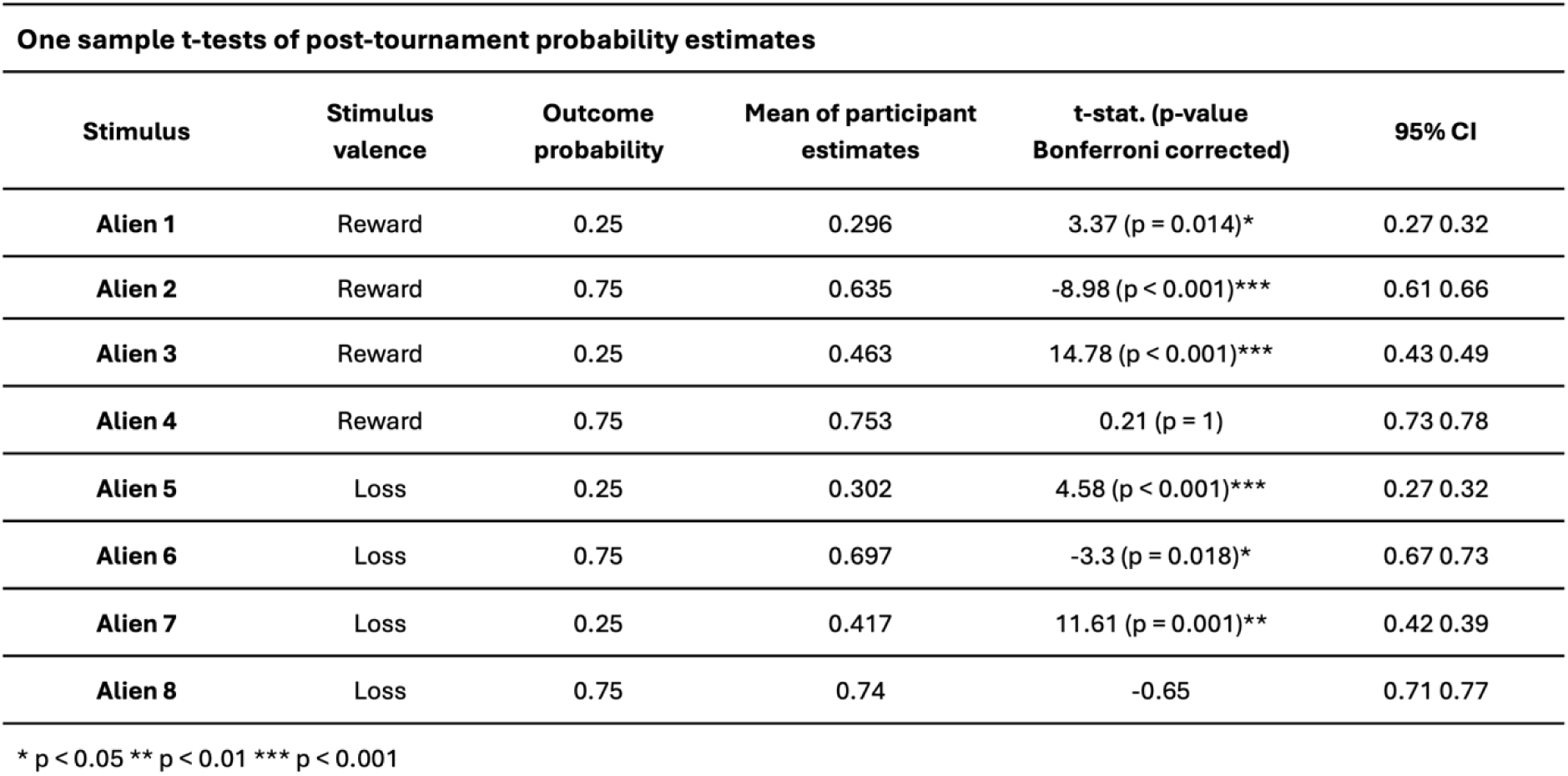
Summary of Bonferroni corrected one-sample t-tests comparing participants’ post-tournament probability estimates against each of the stimuli’s true probability. The average of participants’ post-tournament probability estimates for each stimulus demonstrates that participants tended to approximate the true outcome probabilities. Participants were most accurate in estimating the two stimuli with the highest absolute expected values (Alien 4 and Alien 8). Note that Bonferroni correction is applied by multiplying the p-value of each comparison by the number of comparisons, these adjusted p-values are then compared to the original significance threshold of 0.05. Asterisks therefore indicate whether a one sample t-test was significant at this threshold.

**Supplementary Figure 3:**
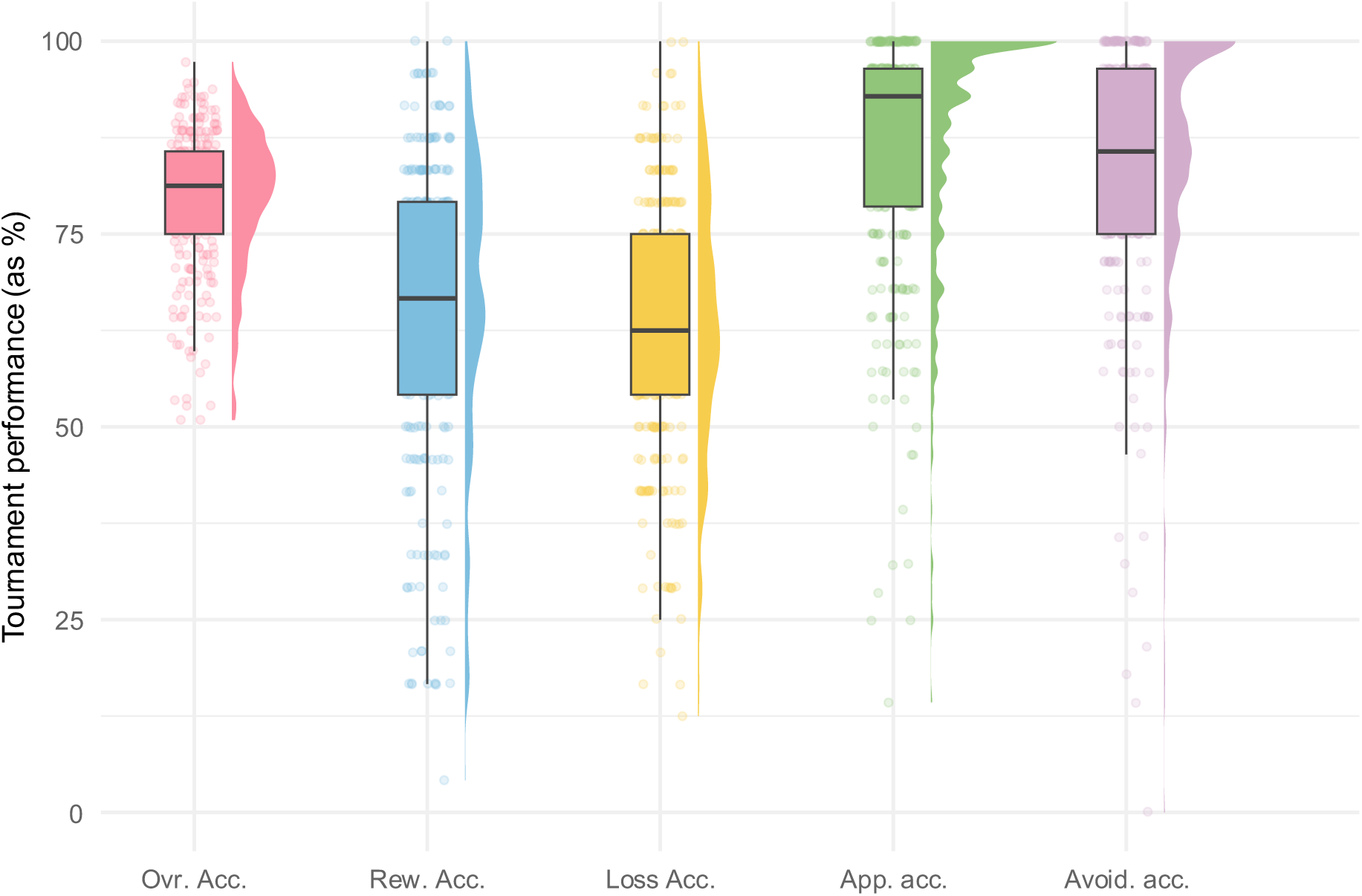
Rain cloud plots visualising participants’ performance in the tournament task. **From left to right: Overall accuracy** the percentage of trials in which participant chose the correct option, where the correct option was defined as the option which had the highest expected value (outcome magnitude * outcome probability); **Reward accuracy** the percentage of trials where participants chose the correct option when both options were stimuli associated with reward; **Loss accuracy** the percentage of trials where participants chose the correct option when both options were stimuli associated with loss; **Approach accuracy** the percentage of trials where participants chose the best stimulus (i.e. the stimulus with the overall highest expected value in the stimulus set) when it was present in a choice pair; **Avoid accuracy** the percentage of trials where participants chose any stimulus other than the worst stimulus (i.e. the stimulus with the lowest expected value in the stimulus set).

**Table S2:**
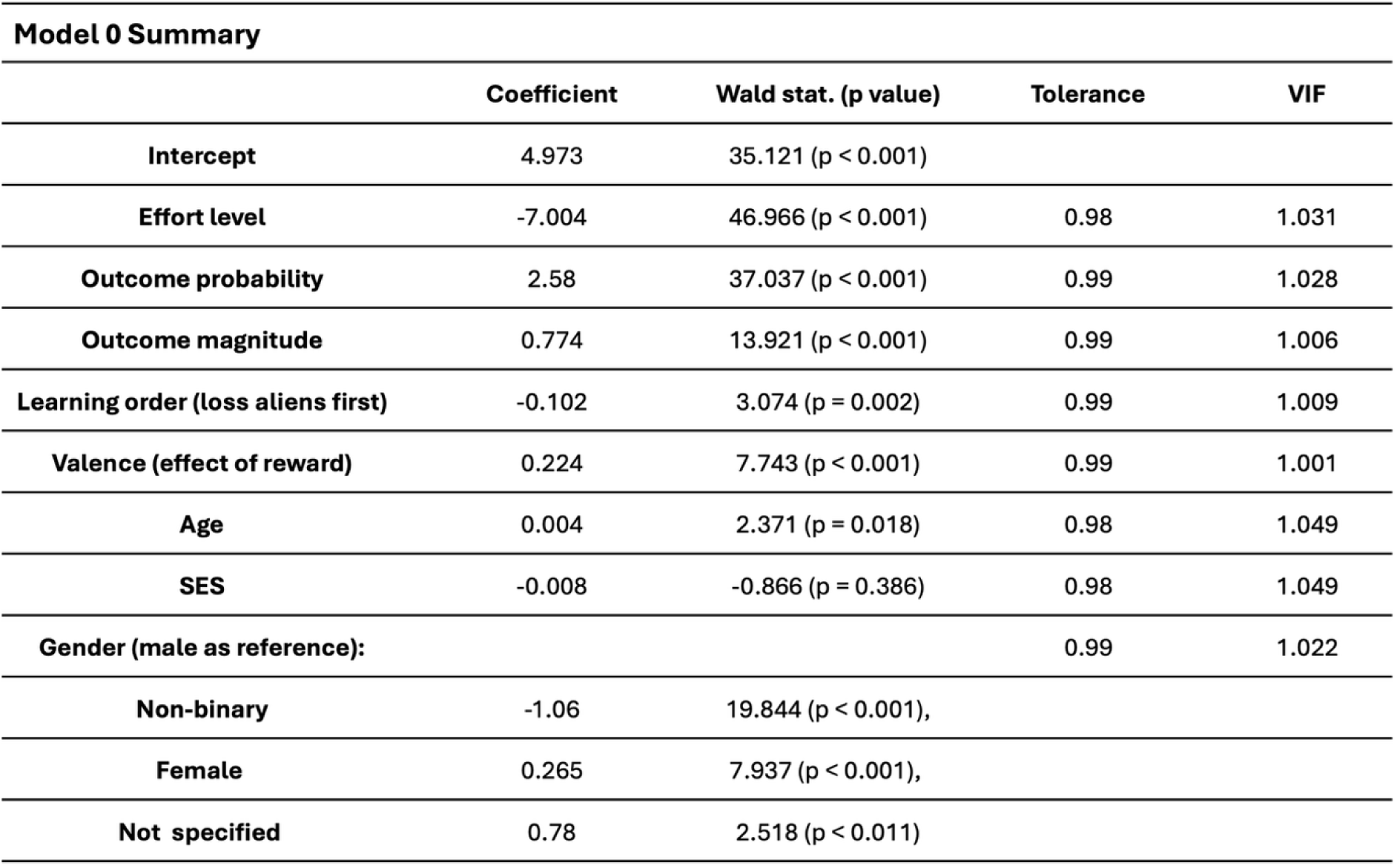
Summary of the logistic regression model (M0) fitted to participants’ trial-by-trial effort choices. The overall model was significant (X^2^ = 4248.173, p < 0.001) and effort level, outcome probability, outcome magnitude, learning order, valence and gender were significant predictors.

**Table S3.**
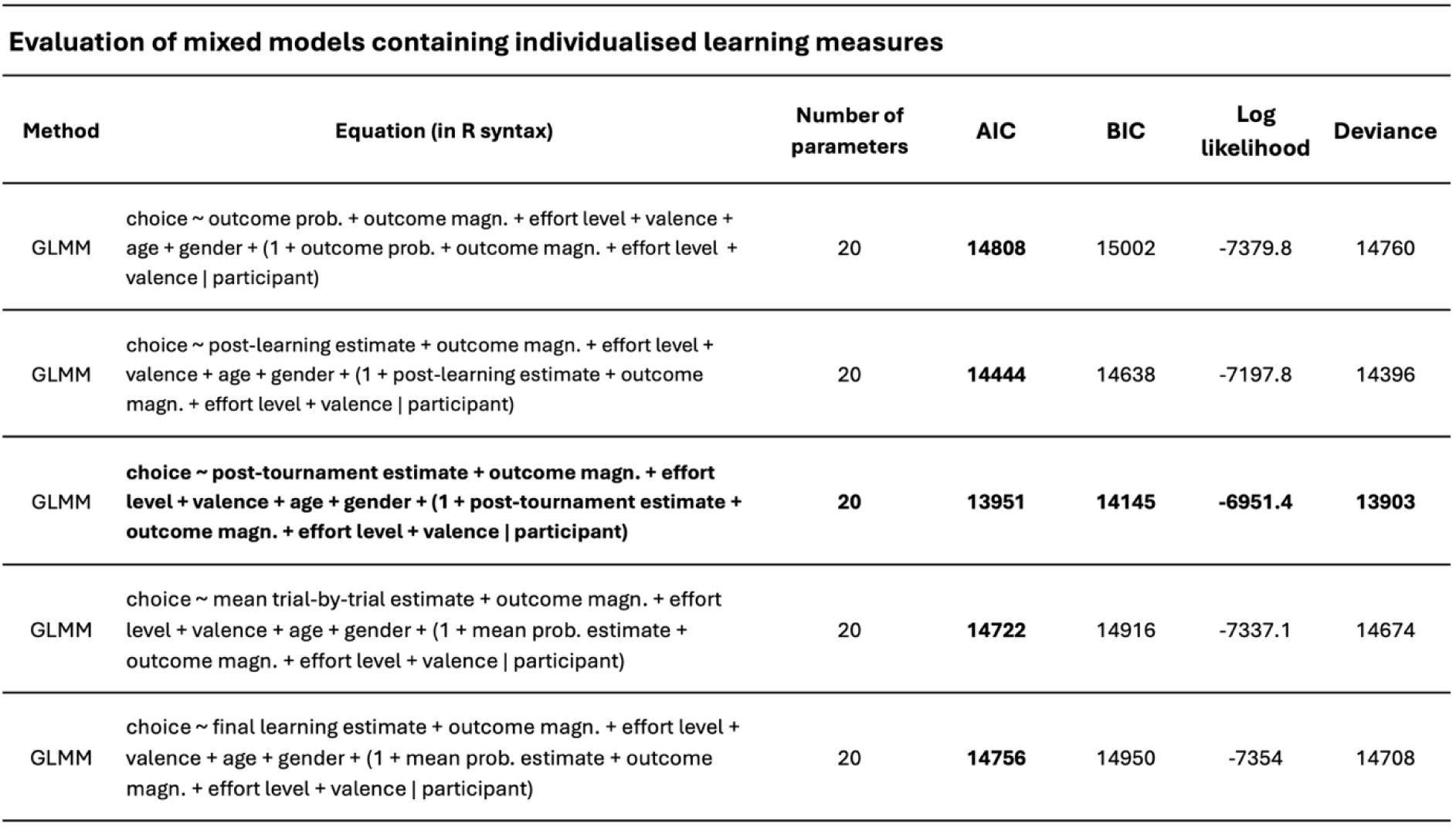
Comparison of the models hich compare the predictive value of individual learning measures. . Table includes: model type, model equation in R (lme4) syntax, number of parameters, AIC, BIC, log likelihood and deviance scores for each model. These models were fit using *lme4* in R (R version 4.4.1; R Core Team, 2024; https://www.R-project.org/) package *lme4 (*version 1.1-35.5; Bates et al. 2024). Details of the individualised learning measures can be found in the **Methods** section. Of these models, Model 3, which included participants’ post-tournament probability estimates, was the best-fitting of all the individualised learning measures based on all model comparison metrics. Importantly, all models which contained an individualised learning metric outperformed the model (Model 1) which contained objective outcome probabilities.

### Correlation amongst individualised learning measures

**Supplementary Figure 4:**
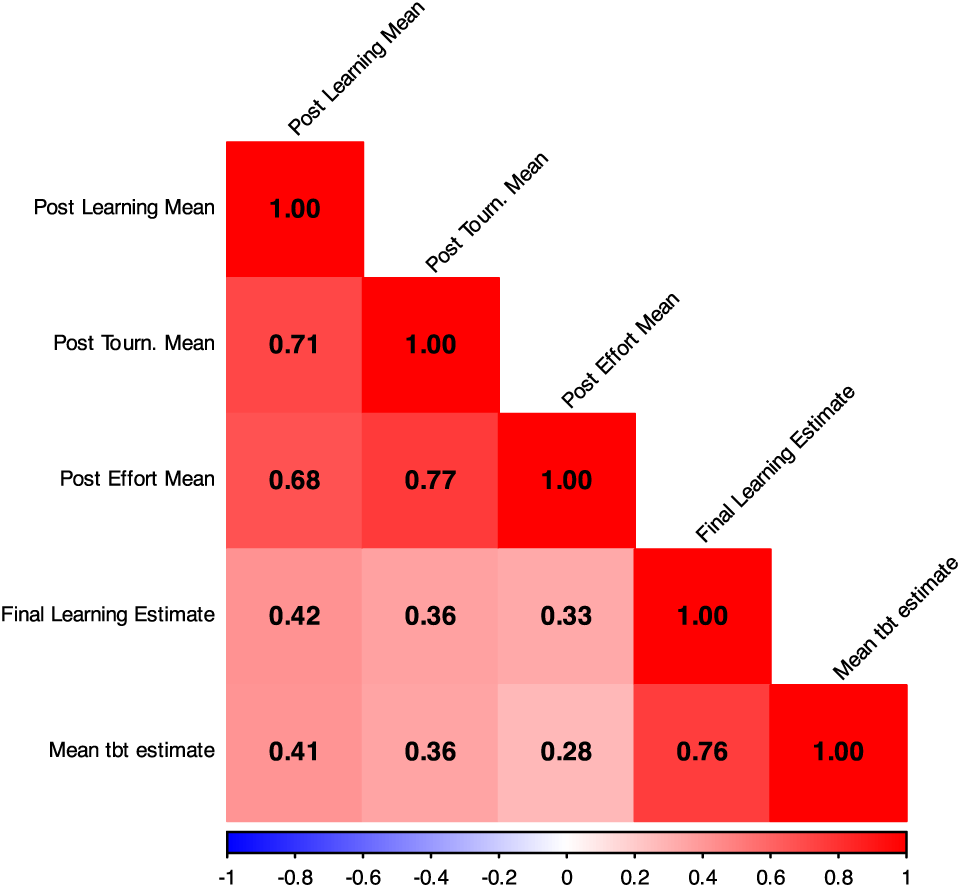
A correlation matrix of the individual learning measures. Correlations amongst all five individualised learning measures used as substitutes for objective outcome probabilities in the generalised linear mixed effects models (GLMMs). All post-task probability estimates were at least moderately correlated with each other. These estimates were less strongly related to measures taken during the learning task (Final Learning Estimate and Mean tbt estimate), but these two measures were strongly associated with one another.

**Supplementary Figure 5:**
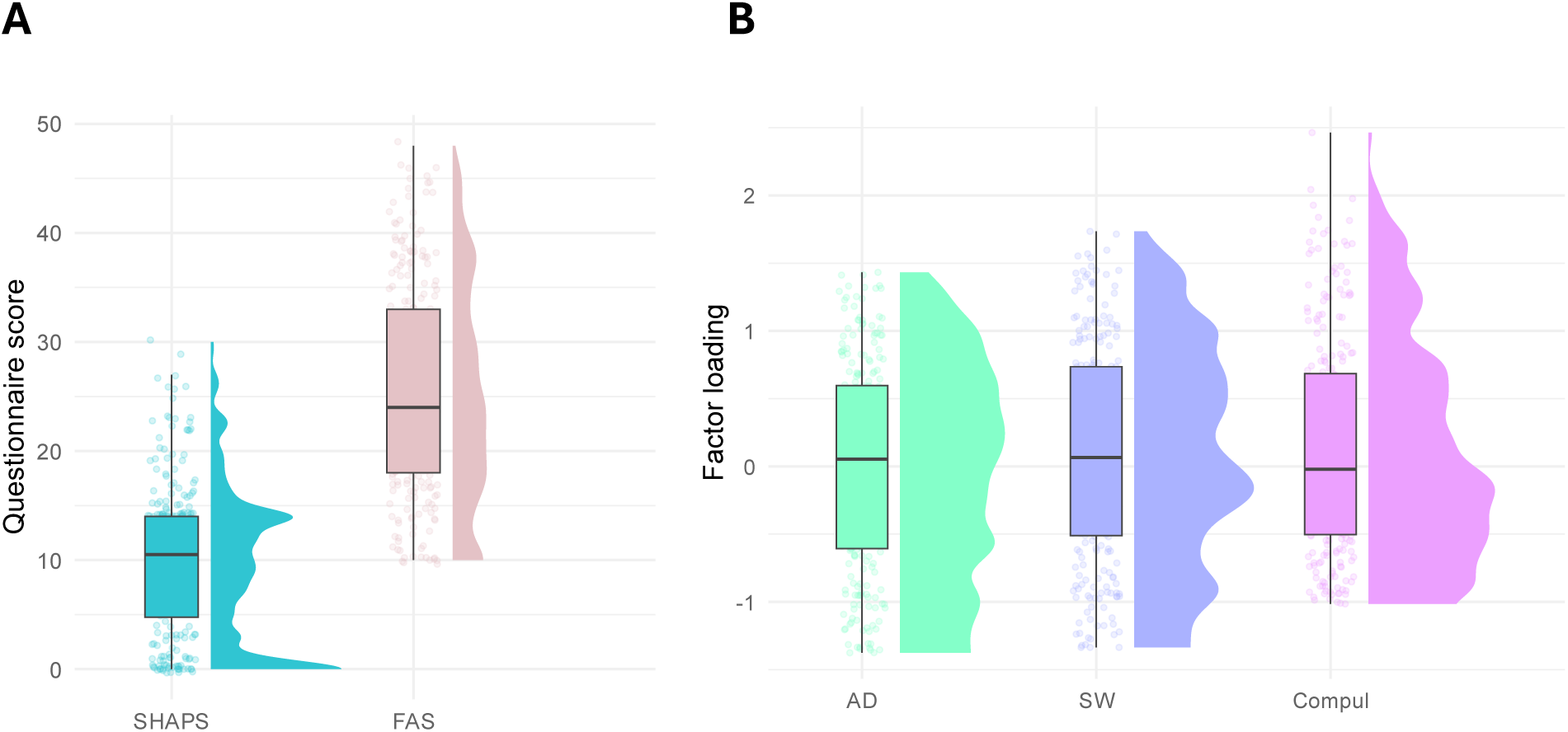
Anhedonia, fatigue scores and transdiagnostic factors. **(A)** Raincloud plots of participants’ summed scores on the SHAPS, a measure of anhedonia and the FAS, a measure of fatigue. **(B)** Raincloud plots of participants’ loadings onto three transdiagnostic mental health factors, anxious-depression, social withdrawal and compulsivity (Gillan et al., 2016) using a reduced set of questionnaire items (Hopkins et al., 2022).

**Table S4:**
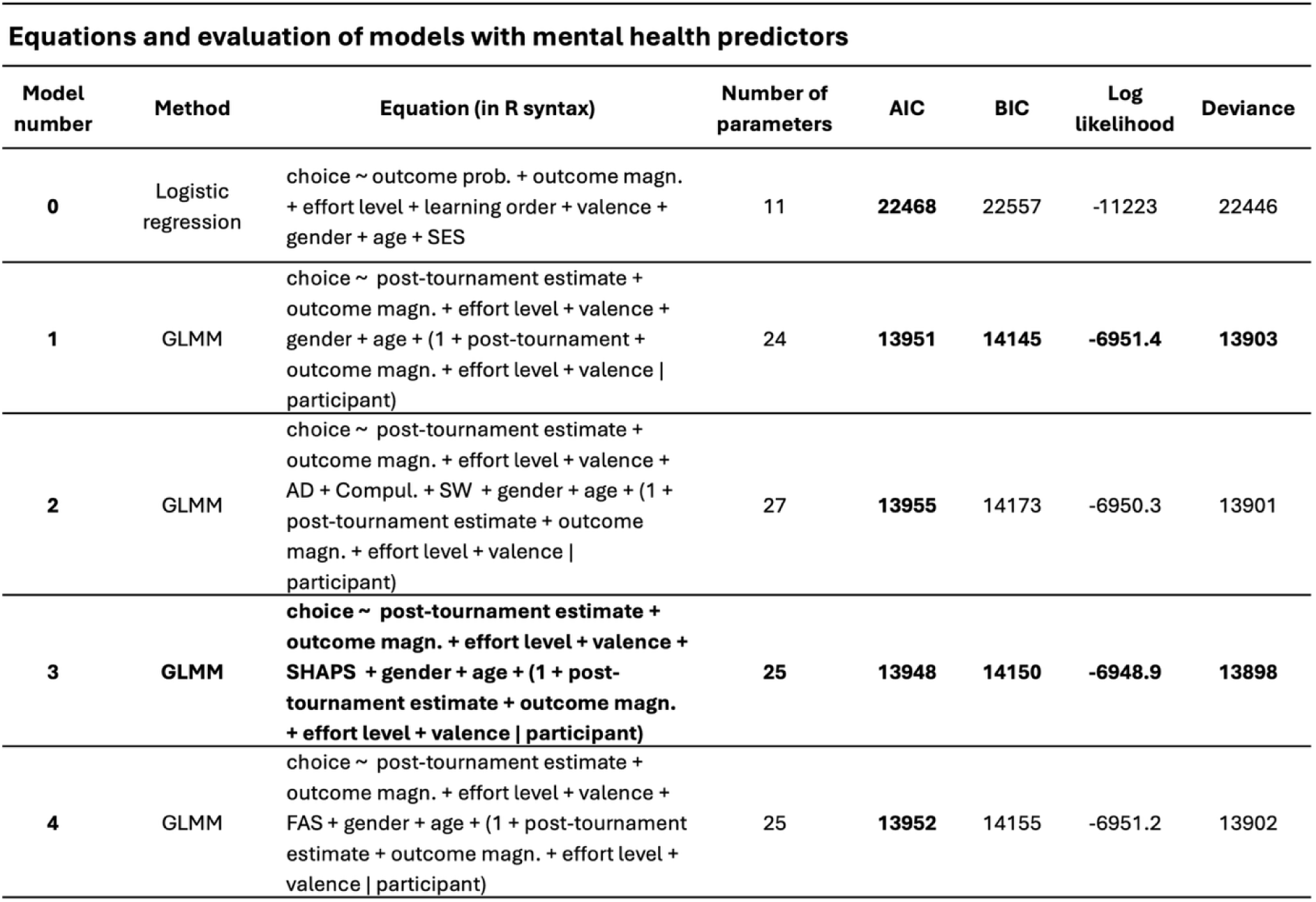
A summary of the models fitted to predict participants trial-by-trial choice behaviour. Including: fitting method used, the model equation written in the syntax of *lme4* in R (R version 4.4.1; R Core Team, 2024; https://www.R-project.org/) package *lme4 (*version 1.1-35.5; Bates et al. 2024), the total number of model parameters, model AIC and BIC scores, model log likelihood and model deviance scores. Here we compare the performance of the three models which include mental health predictors, against the performance of a logistic regression (M0) and the winning model that did not include a mental health predictor (M1). Over and above Model 0 and Model 1, Model 3, which included the SHAPS as a fixed effect predictor, best fit participants’ trial-by-trial choice data on all model comparison measures except BIC, which punishes more severely for additional model complexity.

### Exploratory analysis of the interaction between anhedonia and decision variables

As our primary analysis identified SHAPS scores as a significant predictor of trial-by-trial effort choice, we conducted further analyses to explore whether SHAPS scores significantly interacted with participants’ probability estimates, stimulus outcome magnitude, effort level or valence. For each interaction, we fit a separate exploratory model (EM) based on the winning model (M3):

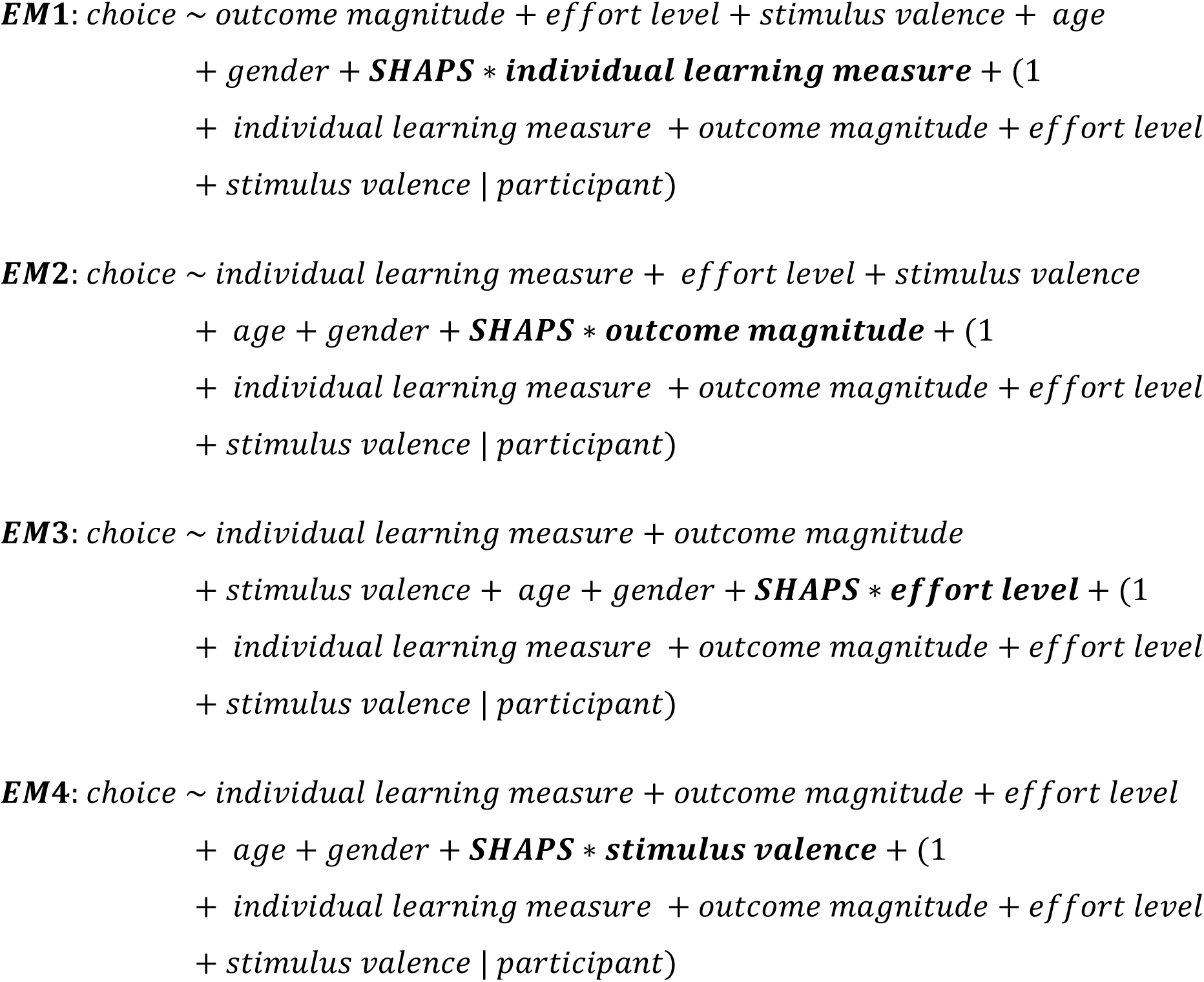

**Supplementary Figure 6:**
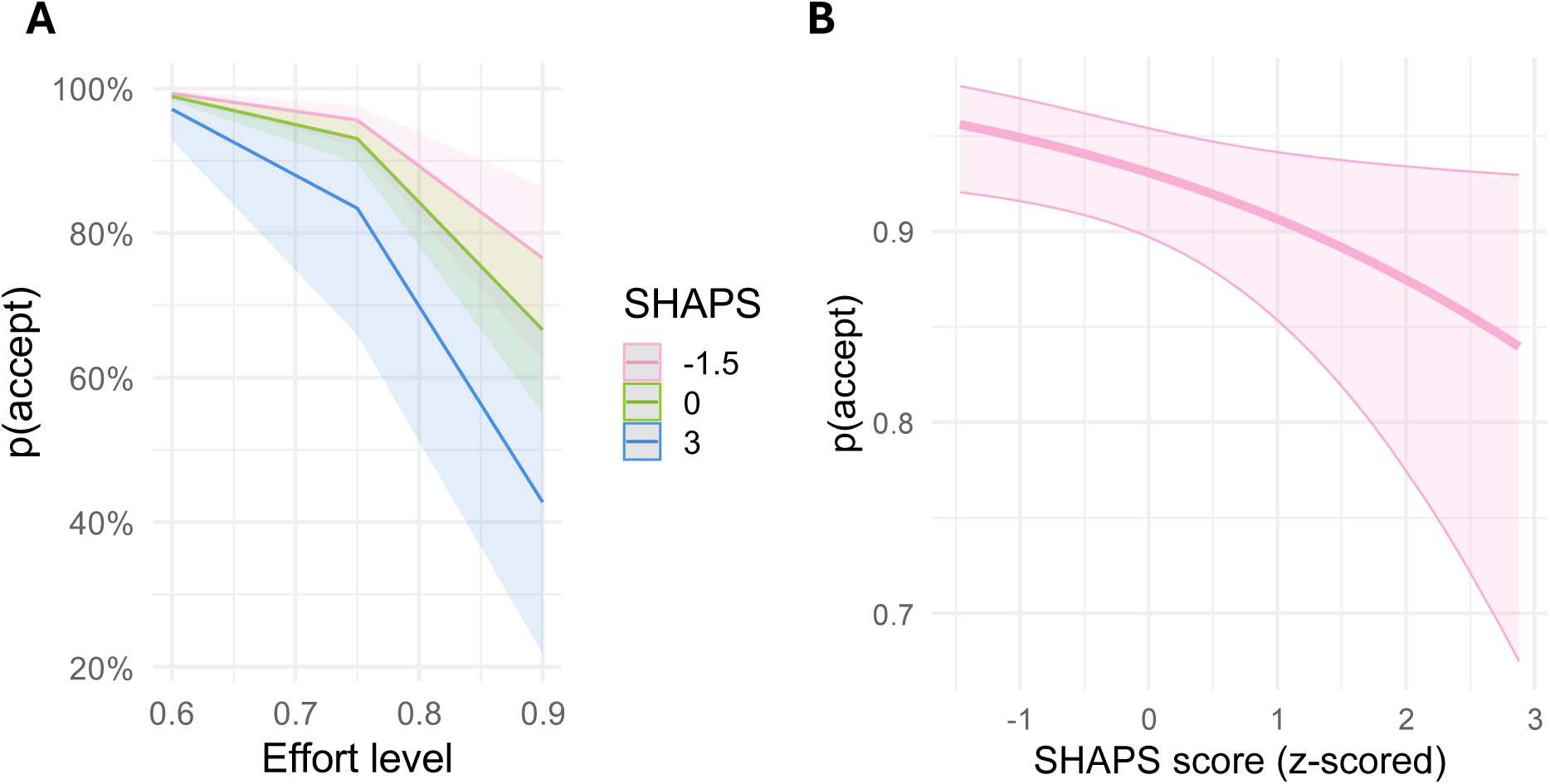
Line plots of the probability of accepting and SHAPS scores. **(A)** A line plot representing predictions from the winning model (Model 3) of the effect of SHAPS scores (low, average and high scorers on the SHAPS) on the probability of accepting across effort levels. Please note this is only for visualisation purposes. Shaded areas are 95% CIs. **(B)** A line plot representing the fixed effect of SHAPS scores on the probability of accepting an effort offer. Shaded areas are 95% CIs.

**Table S5:**
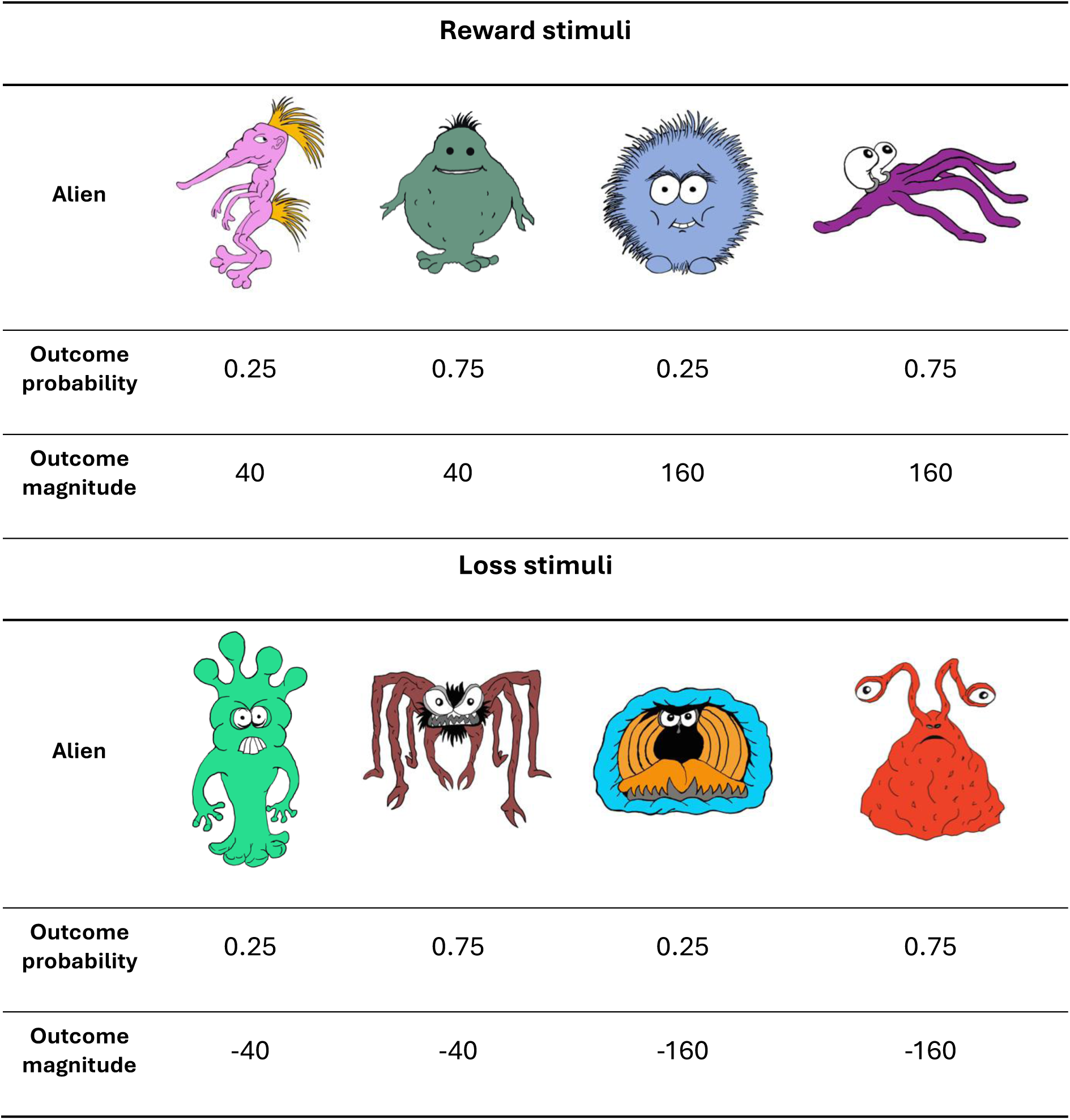
The eight aliens used for the learning, tournament and effort decision-making task. Each alien is presented along with their respective outcome probabilities and outcome magnitude. Each stimulus was associated with a unique combination of outcome magnitude and outcome probability.

## References

1. Hull, C. L. Principles of Behavior, an Introduction to Behavior Theory. (London: Appleton-Century Company, New York, NY, 1943).

2. Zipf, G. K. Human Behavior and the Principle of Least Effort: An Introduction to Human Ecology. (Ravenio Books, 1949).

3. Pessiglione, M., Vinckier, F., Bouret, S., Daunizeau, J. & Le Bouc, R. Why not try harder? Computational approach to motivation deficits in neuro-psychiatric diseases. Brain 141, 629–650 (2018).

4. Westbrook, A., Kester, D. & Braver, T. S. What Is the Subjective Cost of Cognitive Effort? Load, Trait, and Aging Effects Revealed by Economic Preference. PLOS ONE 8, e68210 (2013).

5. Wu, R., Ferguson, A. M. & Inzlicht, M. Do humans prefer cognitive effort over doing nothing? Journal of Experimental Psychology: General 152, 1069–1079 (2023).

6. Husain, M. & Roiser, J. P. Neuroscience of apathy and anhedonia: a transdiagnostic approach. Nat Rev Neurosci 19, 470–484 (2018).

7. Bonnelle, V. et al. Characterization of reward and effort mechanisms in apathy. Journal of Physiology-Paris 109, 16–26 (2015).

8. Schmidt, L., Palminteri, S., Lafargue, G. & Pessiglione, M. Splitting Motivation: Unilateral Effects of Subliminal Incentives. Psychol Sci 21, 977–983 (2010).

9. Jurgelis, M. et al. Heightened effort discounting is a common feature of both apathy and fatigue. Sci Rep 11, 22283 (2021).

10. Mehrhof, S. Z. & Nord, C. L. A common alteration in effort-based decision-making in apathy, anhedonia, and late circadian rhythm. eLife 13, (2024).

11. Cléry-Melin, M.-L. et al. Why Don’t You Try Harder? An Investigation of Effort Production in Major Depression. PLOS ONE 6, e23178 (2011).

12. Klein-Flügge, M. C., Kennerley, S. W., Saraiva, A. C., Penny, W. D. & Bestmann, S. Behavioral Modeling of Human Choices Reveals Dissociable Effects of Physical Effort and Temporal Delay on Reward Devaluation. PLOS Computational Biology 11, e1004116 (2015).

13. Meyniel, F. et al. A specific role for serotonin in overcoming effort cost. eLife 5, e17282 (2016).

14. Müller, T., Klein-Flügge, M. C., Manohar, S. G., Husain, M. & Apps, M. A. J. Neural and computational mechanisms of momentary fatigue and persistence in effort-based choice. Nat Commun 12, 4593 (2021).

15. Treadway, M. T., Buckholtz, J. W., Schwartzman, A. N., Lambert, W. E. & Zald, D. H. Worth the ‘EEfRT’? The Effort Expenditure for Rewards Task as an Objective Measure of Motivation and Anhedonia. PLOS ONE 4, e6598 (2009).

16. Treadway, M. T., Bossaller, N. A., Shelton, R. C. & Zald, D. H. Effort-based decision-making in major depressive disorder: A translational model of motivational anhedonia. Journal of Abnormal Psychology 121, 553–558 (2012).

17. Kuhn, M. et al. Computational Phenotyping of Effort-Based Decision Making in Unmedicated Adults with Remitted Depression. Biological Psychiatry: Cognitive Neuroscience and Neuroimaging (2025) doi:10.1016/j.bpsc.2025.02.006.

18. Browning, M., Behrens, T. E., Jocham, G., O’Reilly, J. X. & Bishop, S. J. Anxious individuals have difficulty learning the causal statistics of aversive environments. Nat Neurosci 18, 590–596 (2015).

19. Dubois, M. & Hauser, T. U. Value-free random exploration is linked to impulsivity. Nat Commun 13, 4542 (2022).

20. Gibbs-Dean, T. et al. Belief updating in psychosis, depression and anxiety disorders: A systematic review across computational modelling approaches. Neuroscience & Biobehavioral Reviews 147, 105087 (2023).

21. Pulcu, E. & Browning, M. The Misestimation of Uncertainty in Affective Disorders. Trends in Cognitive Sciences 23, 865–875 (2019).

22. Sandhu, T. R., Xiao, B. & Lawson, R. P. Transdiagnostic computations of uncertainty: towards a new lens on intolerance of uncertainty. Neuroscience & Biobehavioral Reviews 148, 105123 (2023).

23. Rouhani, N., Norman, K. A. & Niv, Y. Dissociable effects of surprising rewards on learning and memory. J Exp Psychol Learn Mem Cogn 44, 1430–1443 (2018).

24. Rouhani, N. & Niv, Y. Signed and unsigned reward prediction errors dynamically enhance learning and memory. eLife 10, e61077 (2021).

25. Frank, M. J., Seeberger, L. C. & O’Reilly, R. C. By Carrot or by Stick: Cognitive Reinforcement Learning in Parkinsonism. Science 306, 1940–1943 (2004).

26. Lawson, R. P. et al. The habenula encodes negative motivational value associated with primary punishment in humans. Proceedings of the National Academy of Sciences 111, 11858–11863 (2014).

27. Lawson, R. P. et al. Disrupted habenula function in major depression. Mol Psychiatry 22, 202–208 (2017).

28. Kessler, R. C. et al. Anxious and non-anxious major depressive disorder in the World Health Organization World Mental Health Surveys. Epidemiology and Psychiatric Sciences 24, 210–226 (2015).

29. Hezemans, F. H., Wolpe, N. & Rowe, J. B. Apathy is associated with reduced precision of prior beliefs about action outcomes. Journal of Experimental Psychology: General 149, 1767 (2020).

30. Nord, C. L. et al. Vigour in active avoidance. Sci Rep 7, 60 (2017).

31. Stein, M. B. & Paulus, M. P. Imbalance of Approach and Avoidance: The Yin and Yang of Anxiety Disorders. Biological Psychiatry 66, 1072–1074 (2009).

32. Palminteri, S., Lefebvre, G., Kilford, E. J. & Blakemore, S.-J. Confirmation bias in human reinforcement learning: Evidence from counterfactual feedback processing. PLOS Computational Biology 13, e1005684 (2017).

33. Kahneman, D. & Tversky, A. Prospect Theory: An Analysis of Decision Under Risk. in World Scientific Handbook in Financial Economics Series vol. 4 99–127 (WORLD SCIENTIFIC, 2013).

34. Rzepa, E., Fisk, J. & McCabe, C. Blunted neural response to anticipation, effort and consummation of reward and aversion in adolescents with depression symptomatology. J Psychopharmacol 31, 303–311 (2017).

35. Lockwood, P. L. et al. Prosocial apathy for helping others when effort is required. Nat Hum Behav 1, 0131 (2017).

36. Bishop, S. J. & Gagne, C. Anxiety, Depression, and Decision Making: A Computational Perspective. Annual Review of Neuroscience 41, 371–388 (2018).

37. Huang, H., Thompson, W. & Paulus, M. P. Computational Dysfunctions in Anxiety: Failure to Differentiate Signal From Noise. Biological Psychiatry 82, 440–446 (2017).

38. Wise, T. & Dolan, R. J. Associations between aversive learning processes and transdiagnostic psychiatric symptoms in a general population sample. Nat Commun 11, 4179 (2020).

39. Bedder, R. L., Pisupati, S. & Niv, Y. Modelling Rumination as a State-Inference Process. Proceedings of the Annual Meeting of the Cognitive Science Society 45, (2023).

40. Bustamante, L. A. et al. Major depression symptom severity associations with willingness to exert effort and patch foraging strategy. Psychological Medicine 54, 4396–4407 (2024).

41. Gagne, C., Agai, S., Ramiro, C., Dayan, P. & Bishop, S. Biased belief priors versus biased belief updating: Differential correlates of depression and anxiety. PLOS Computational Biology 18, e1010176 (2022).

42. Lynch, C. J., Gunning, F. M. & Liston, C. Causes and Consequences of Diagnostic Heterogeneity in Depression: Paths to Discovering Novel Biological Depression Subtypes. Biological Psychiatry 88, 83–94 (2020).

43. Robinson, O. J. & Chase, H. W. Learning and Choice in Mood Disorders: Searching for the Computational Parameters of Anhedonia. Computational Psychiatry 1, 208–233 (2017).

44. Caspi, A. et al. The p Factor: One General Psychopathology Factor in the Structure of Psychiatric Disorders? Clinical Psychological Science 2, 119–137 (2014).

45. Cuthbert, B. N. & Insel, T. R. Toward the future of psychiatric diagnosis: the seven pillars of RDoC. BMC Med 11, 126 (2013).

46. Gillan, C. M., Kosinski, M., Whelan, R., Phelps, E. A. & Daw, N. D. Characterizing a psychiatric symptom dimension related to deficits in goal-directed control. eLife 5, e11305 (2016).

47. Patzelt, E. H., Kool, W., Millner, A. J. & Gershman, S. J. The transdiagnostic structure of mental effort avoidance. Sci Rep 9, 1689 (2019).

48. Ang, Y.-S., Gelda, S. E. & Pizzagalli, D. A. Cognitive effort-based decision-making in major depressive disorder. Psychological Medicine 53, 4228–4235 (2023).

49. Wise, T., Robinson, O. J. & Gillan, C. M. Identifying Transdiagnostic Mechanisms in Mental Health Using Computational Factor Modeling. Biological Psychiatry 93, 690–703 (2023).

50. Foa, E. B. et al. The Obsessive-Compulsive Inventory: Development and validation of a short version. Psychological Assessment 14, 485–496 (2002).

51. Marin, R. S., Biedrzycki, R. C. & Firinciogullari, S. Reliability and validity of the apathy evaluation scale. Psychiatry Research 38, 143–162 (1991).

52. Liebowitz, M. R. Social Phobia. (1987) doi:10.1159/000414022.

53. Zung, W. W. K. A Self-Rating Depression Scale. Archives of General Psychiatry 12, 63–70 (1965).

54. Spielberger, C., Gorsuch, R., Lushene, R., Vagg, P. & Jacobs, G. Manual for the State-Trait Anxiety Inventory. (Consulting Psychologists Press., 1983).

55. Valton, V. et al. A computational approach to understanding effort-based decision-making in depression. bioRxiv 2024.06.17.599286 (2024) doi:10.1101/2024.06.17.599286.

56. Müller, T., Milton, J., Husain, M. & Apps, M. A. Computational signatures of exertion and rest underlie moment-to-moment dynamics of subjective perceptions of effort and fatigue. (2025).

57. Gagne, C., Zika, O., Dayan, P. & Bishop, S. J. Impaired adaptation of learning to contingency volatility in internalizing psychopathology. eLife 9, e61387 (2020).

58. Huys, Q. J., Pizzagalli, D. A., Bogdan, R. & Dayan, P. Mapping anhedonia onto reinforcement learning: a behavioural meta-analysis. Biol Mood Anxiety Disord 3, 12 (2013).

59. Hopkins, A. K., Gillan, C., Roiser, J., Wise, T. & Sidarus, N. Optimising the measurement of anxious-depressive, compulsivity and intrusive thought and social withdrawal transdiagnostic symptom dimensions. Preprint at 10.31234/osf.io/q83sh (2022).

60. Snaith, R. P. et al. A scale for the assessment of hedonic tone the Snaith-Hamilton Pleasure Scale. Br J Psychiatry 167, 99–103 (1995).

61. Franken, I. H. A., Rassin, E. & Muris, P. The assessment of anhedonia in clinical and non-clinical populations: Further validation of the Snaith–Hamilton Pleasure Scale (SHAPS). Journal of Affective Disorders 99, 83–89 (2007).

62. Shahid, A., Wilkinson, K., Marcu, S. & Shapiro, C. M. Fatigue Assessment Scale (FAS). in STOP, THAT and One Hundred Other Sleep Scales (eds Shahid, A., Wilkinson, K., Marcu, S. & Shapiro, C. M.) 161–162 (Springer New York, New York, NY, 2011). doi:10.1007/978-1-4419-9893-4_33.

