## Supplementary Materials for "Prior learning shapes effort choice, revealing valence asymmetries and disruption in anhedonia"

**Word count (excluding abstract, methods, tables/figures and references):** 5000

**Author note**

Correspondence concerning this manuscript should be addressed to Calum Guinea, Department of Psychology, Downing Place, University of Cambridge, CB2 3EB, United Kingdom. **Email:**. All authors declare no conflict of interest. This study was not preregistered. **Data availability:** The data from this study are available from the corresponding author upon reasonable request. **Funding statement:** CG is funded by the Harding Distinguished Postgraduate Scholarship. BX is funded by the Trinity Henry Barlow Studentship and the Cambridge International Scholarship. This work was supported by a Wellcome Trust Henry Dale Fellowship awarded to RPL. **Ethics:** This work was approved by the Cambridge Psychology Research Ethics Committee (CPREC), ethics number PRE.2019.110.

**Author ORCIDs**

Calum Guinea: https://orcid.org/0009-0006-9444-9838

Bowen Xiao: https://orcid.org/0000-0002-9627-3454

Rebecca P. Lawson: https://orcid.org/0000-0003-1228-1244

#### Abstract

Goal directed behaviour requires the integration of learned beliefs about the world with decisions about whether an outcome is worth the effort required to obtain it. Although value learning and effort-based choice are central to motivated behaviour, they are typically examined in isolation. Here we introduce a novel behavioural paradigm that directly links probabilistic associative learning with effortful decision making, for the first time allowing trial by trial assessment of how learned beliefs guide willingness to exert effort.

| Summary Table 1 | |
|---|---|
| **Sample size** | 252 |
| **Demographics** | |
| Age, mean (SD) | 38.93 (11.92) |
| Gender, number (%) | |
| Male | 125 (49.6) |
| Female | 125 (49.6) |
| Non-binary | 1 (0.4) |
| Rather not say | 1 (0.4) |
| Ethnicity, number (%) | |
| Asian | 15 (5.95) |
| Black | 13 (4.76) |
| Mixed | 6 (2.38) |
| Other | 2 (0.79) |
| White | 217 (86.11) |
| Socioeconomic status, mean (SD) | 5.05 (1.7) |
| **Transdiagnostic factor loadings** | |
| Anxious-depression, mean (SD) | -0.6 (0.76) |
| Compulsivity and intrusive thought, mean (SD) | 0.14 (0.79) |
| Social withdrawal, mean (SD) | 0.1 (0.79) |
| **Questionnaire measures** | |
| SHAPS, mean sum score (SD) | 10.17 (6.93) |
| FAS, mean sum score (SD) | 25.73 (9.6) |

**Table 1: Demographic information of the sample**. Participant loadings onto the three transdiagnostic factors and scores on self-report measures of anhedonia and fatigue.

$$\boldsymbol{M1}: \ choice \sim \boldsymbol{individual\ learning\ measure} + outcome\ magnitude + effort\ level + stimulus\ valence + age + gender + (1 + \boldsymbol{individual\ learning\ measure} + outcome\ magnitude + effort\ level + stimulus\ valence \mid participant)$$

Lastly, in this general population sample, we were interested in whether participants' effort choices were related to a set of relevant psychiatric dimensions and clinical symptoms (e.g. three transdiagnostic factor scores comprising anxious-depression, compulsivity and intrusive thought and social withdrawal; in addition to anhedonia and fatigue). We therefore fit the following suite of models (bold text indicates where models have changed from the previous model):

$$\boldsymbol{M2}: \ choice \sim individual\ learning\ measure + outcome\ magnitude + effort\ level + stimulus\ valence + age + gender + \boldsymbol{AD} + \boldsymbol{Compul} + \boldsymbol{SW} + (1 + individual\ learning\ measure + outcome\ magnitude + effort\ level + stimulus\ valence \mid participant)$$

$$\boldsymbol{M3}: \ choice \sim individual\ learning\ measure + outcome\ magnitude + effort\ level + stimulus\ valence + age + gender + \boldsymbol{SHAPS} + (1 + individual\ learning\ measure + outcome\ magnitude + effort\ level + stimulus\ valence \mid participant)$$

$$\boldsymbol{M4}: \ choice \sim individual\ learning\ measure + outcome\ magnitude + effort\ level + stimulus\ valence + age + gender + \boldsymbol{FAS} + (1 + individual\ learning\ measure + outcome\ magnitude + effort\ level + stimulus\ valence \mid participant)$$

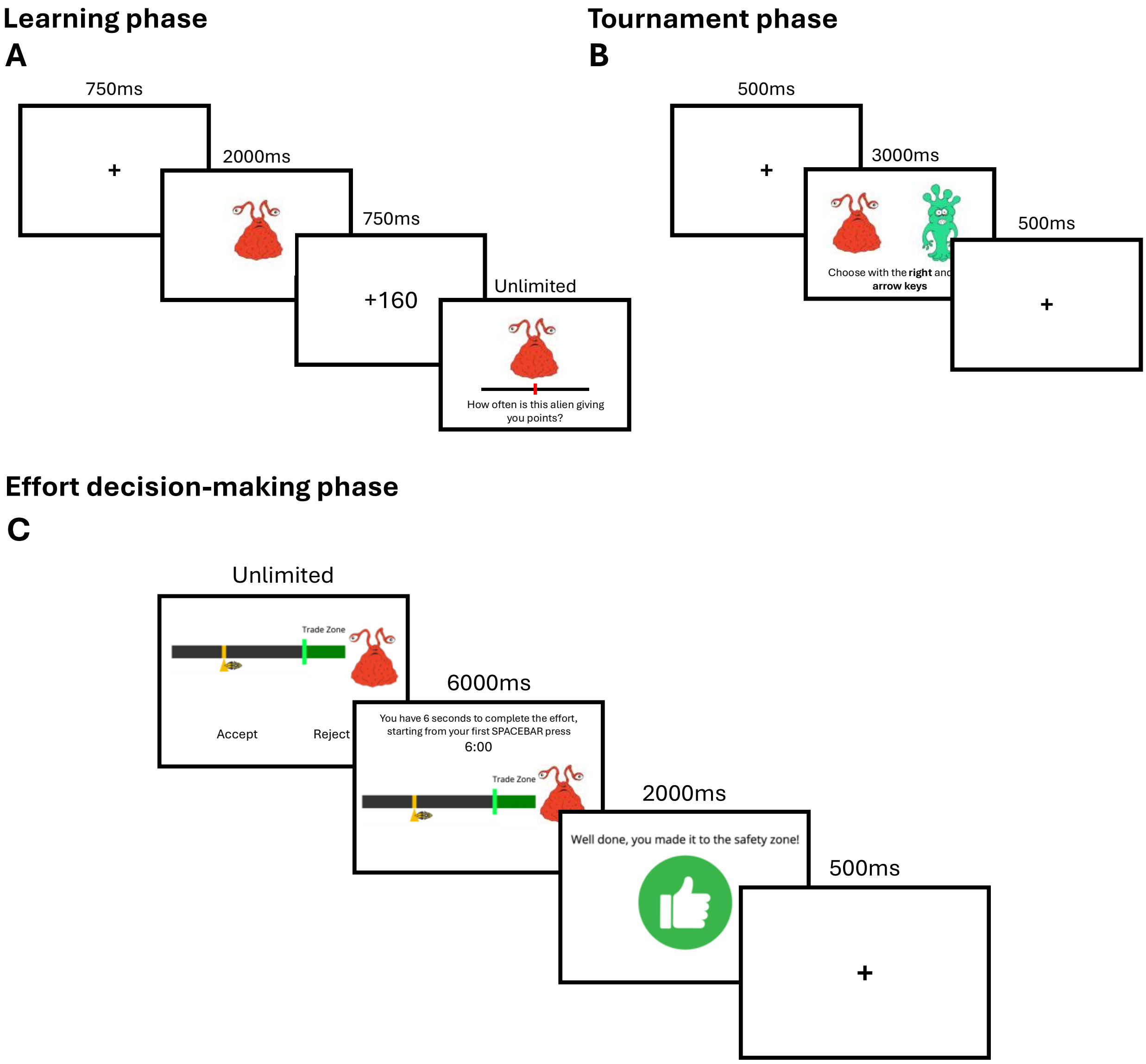


**Figure 1: Learning, tournament and effort-decision making phases of the experiment (A)** An example trial from the learning phase of the experiment. Participants first view a fixation cross, then one of the eight stimuli is shown on screen before the outcome of the trial appears on the next screen. For reward stimuli, when the alien gives points, a positive tone is played when the amount appears on screen and for loss trials, a negative tone is played when the alien takes points. For both alien types, a neutral tone is played when the outcome is zero. At the end of each trial, participants are asked to estimate the percentage of trials that the alien is giving or taking points from them, this estimate is reported using a slider. **(B)** An example trial from the tournament. Participants see a brief fixation cross and are then presented with two of the eight learned stimuli and asked to make a choice between them in 3000ms using the left and right arrow keys on their keyboard. If no choice is made this is recorded as no response. If participants make a choice in under 3000ms the next pair of stimuli appear on the screen after a brief delay. **(C)** An example trial from the effort decision-making phase where the participant accepts the offer and meets/exceeds the effort requirement. On each effort trial, participants are presented with one of the eight stimuli they learned about previously and an effort amount visualised as the distance from a Trade/Safety zone on a horizontal bar. Participants

are required to accept or reject based on the effort required and what they remember about the outcome probability and magnitude associated with the given stimulus. When accepting the offer, participants are taken to a screen which includes a countdown timer that counts down to zero from 6000ms as soon as the participant makes the first spacebar press. Rejecting the effort offer takes participants to a waiting screen that also last for 6000ms.

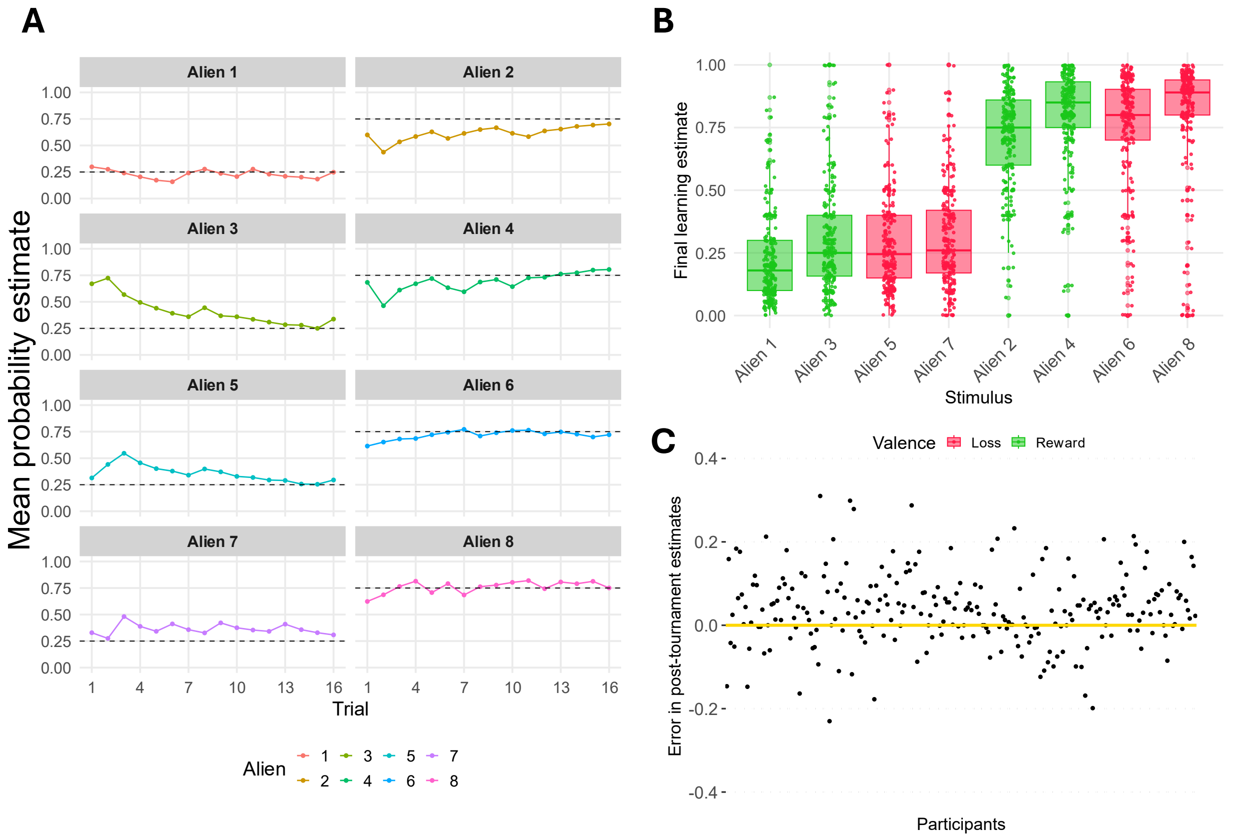


**Figure 2: Summary of performance in the learning phase of the task (A)** The average of all participants' probability estimates across the learning trials. Here we take the mean rating of all participants' estimates for each stimulus on each trial (coloured lines) to visualise the trajectory of participants' learning and how, across trials, their ratings tend toward the true outcome probability (dashed black line). **(B)** Box and scatter plots of participants' final learning estimate for each stimulus. The stimuli have been ordered to group the low probability (p = 0.25) and high probability (p = 0.75) stimuli. Green indicates reward stimuli and red indicates loss stimuli. Participants tended towards the objective outcome probability but there was also notable variability in participants' estimates. **(C)** Here we take the mean post-tournament probability estimate of each participant and subtract the true average outcome probability from it (0.5), such that participants who on average, overestimate the true outcome probabilities are above the yellow line. This average error indicates there is significant variability in learning fidelity across participants.

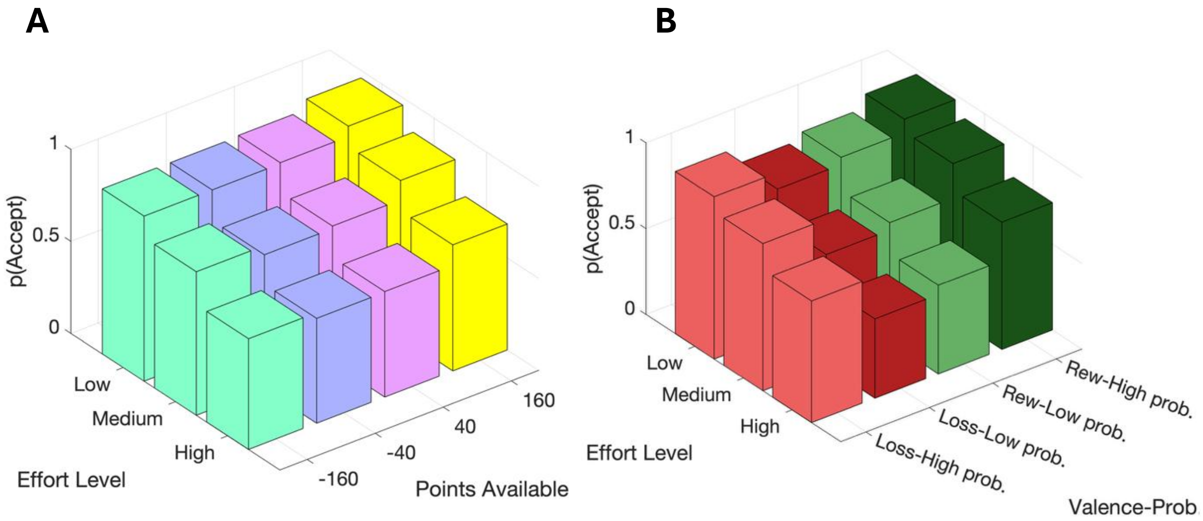


**Figure 3: Probability of accepting as a function of effort level, outcome magnitude and valence-probability. (A)** A 3D bar plot showing how the probability of accepting effort varies as a function of effort level and the amount of points associated with a stimulus. The probability of accepting was highest for the highest outcome magnitude stimuli combined with the lowest effort and lowest for the low magnitude loss stimuli combined with the highest effort level. For each level of outcome magnitude, p(accept) decreased as effort level increased. **(B)** A 3D bar plot showing how the probability of accepting effort varies as a function of effort level and outcome probability for both reward and loss aliens. The probability of accepting was highest for the high probability reward stimuli combined with the lowest effort level and lowest for the low magnitude loss stimuli combined with the highest effort level. For each stimulus valence and outcome probability combination, p(accept) decreased as effort level increased.

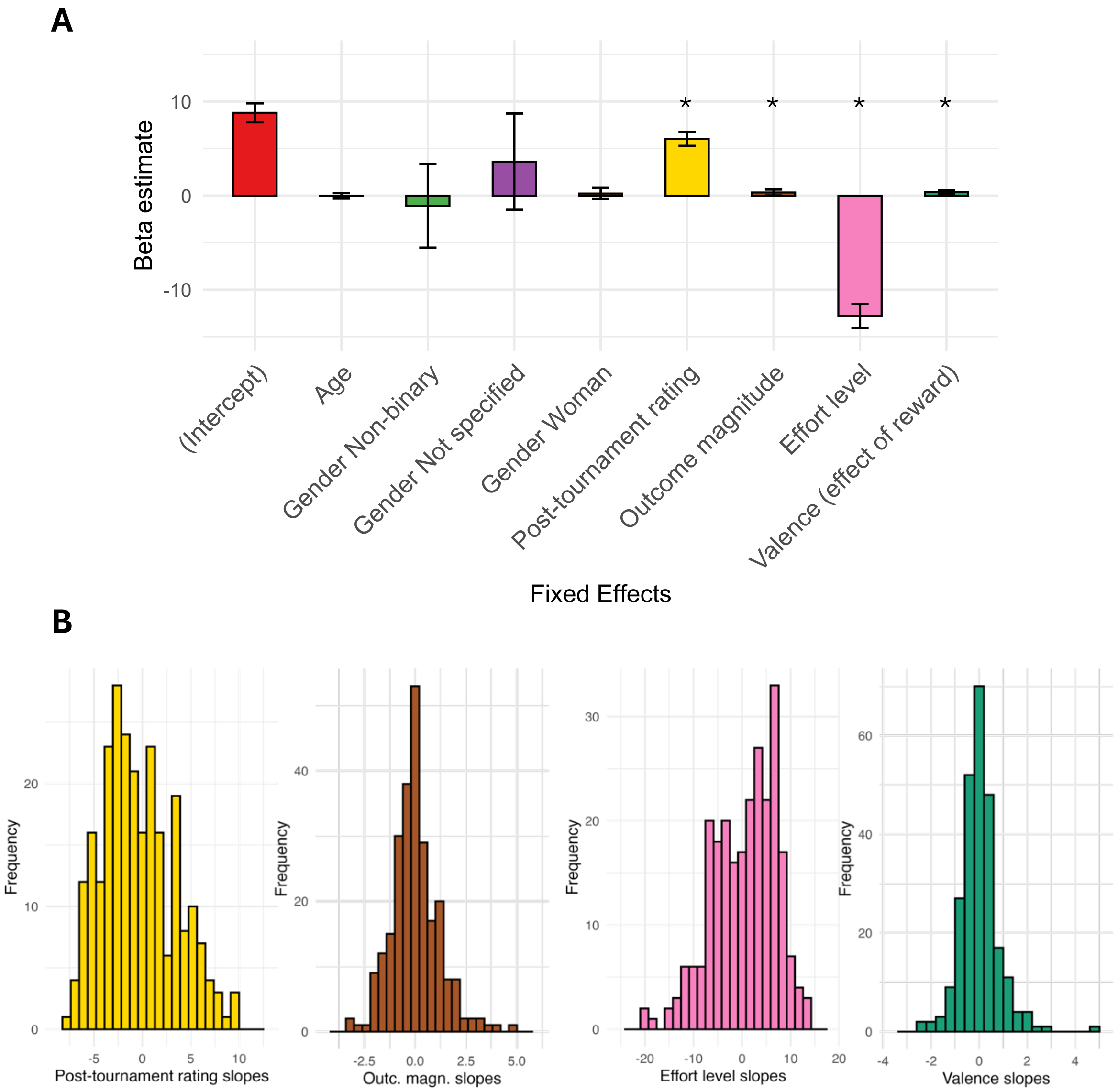


**Figure 4: Bar plot of fixed effect estimates and histograms of the random effects from M1. (A)** Bar plot of the fixed effect estimates from Model 1 (M1), the winning model amongst those fit to compare the performance of individualised learning measures against the objective outcome probabilities. Each bar represents the fixed effect estimate for a predictor from the winning model (M1). Error bars represent the 95% confidence interval estimates for the predictor. Unlike in Model 0 (M0), in the GLMMs, none of the demographic predictors were significantly non-zero predictors of trial-by-trial effort choice. Post-tournament rating, outcome magnitude, effort level and stimulus valence were all significantly non-zero predictors ($p < 0.05$). **(B)** Histograms of the random slope estimates for each of the predictors in the winning model, Model 1 (M1). For example, where the fixed effect estimate for effort level is zero, each participant's slope estimate for the effect of effort level on choice is expressed as a difference from zero. Therefore, participants with a positive slope estimate for effort level are more sensitive to effort level when making effort choices. The slopes for each random effect are approximately normally distributed.

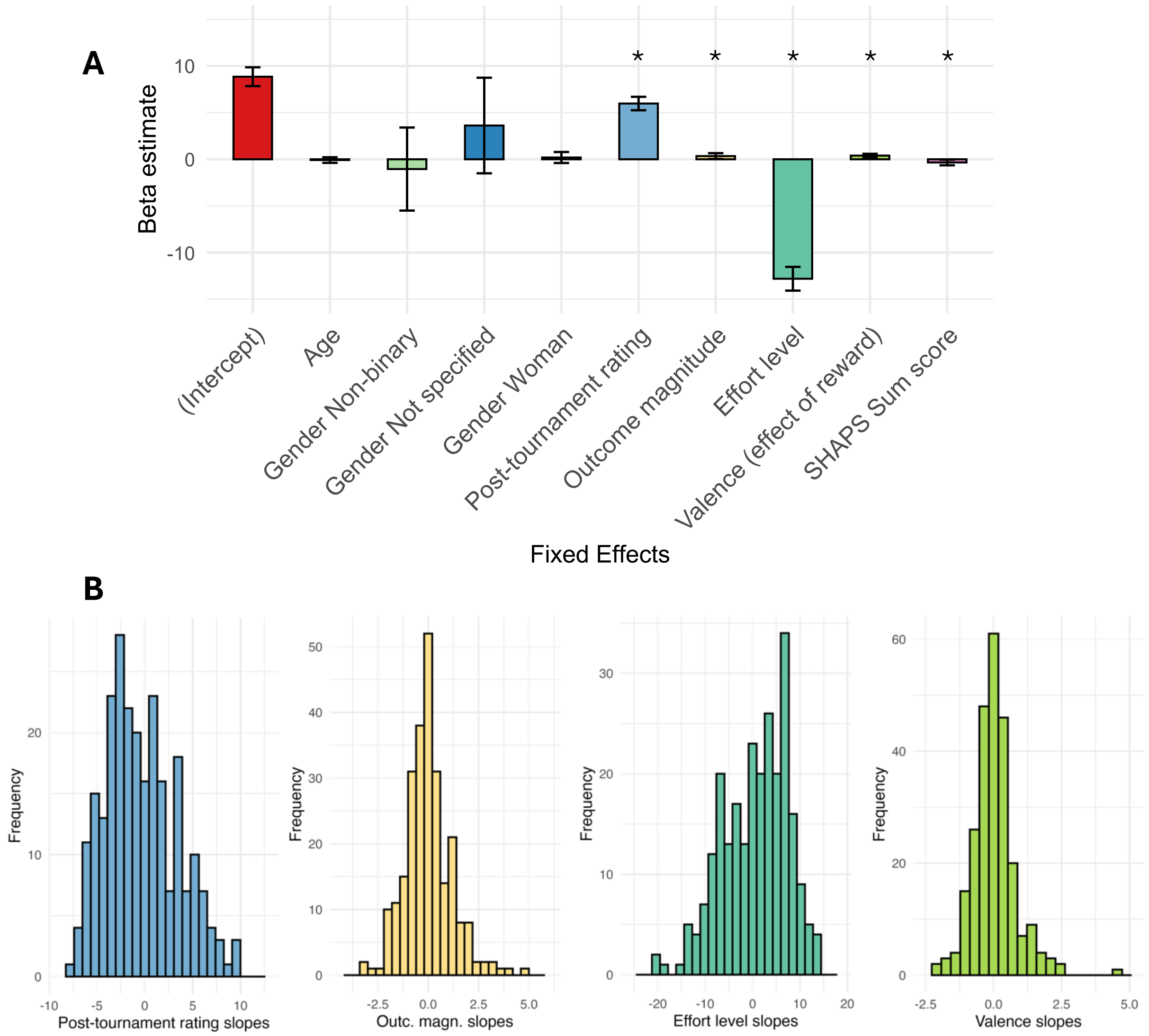


**Figure 5: Bar plot of the fixed effects and histograms of the random effects from M3. (A)** Bar plot of the fixed effect estimates from the overall best-fitting model, which included participants' SHAPS sum scores as a fixed effect predictor (Model 3). Each bar represents the fixed effect estimate for a predictor from the winning GLME model (Model 3), error bars are 95% confidence intervals. All predictors are significant ($p < 0.05$). **(B)** Histograms of the random slopes estimated for each participant from the winning model (Model 4). The slopes fit for each participant are expressed as a difference from zero, where zero is the fixed effect estimate for the predictor. So, for post-tournament rating, a participant with a positive slope estimate suggests that the participant's effort choices are more strongly influenced by their own probability estimate of the stimulus. For each predictor, the slope estimates are approximately normally distributed.

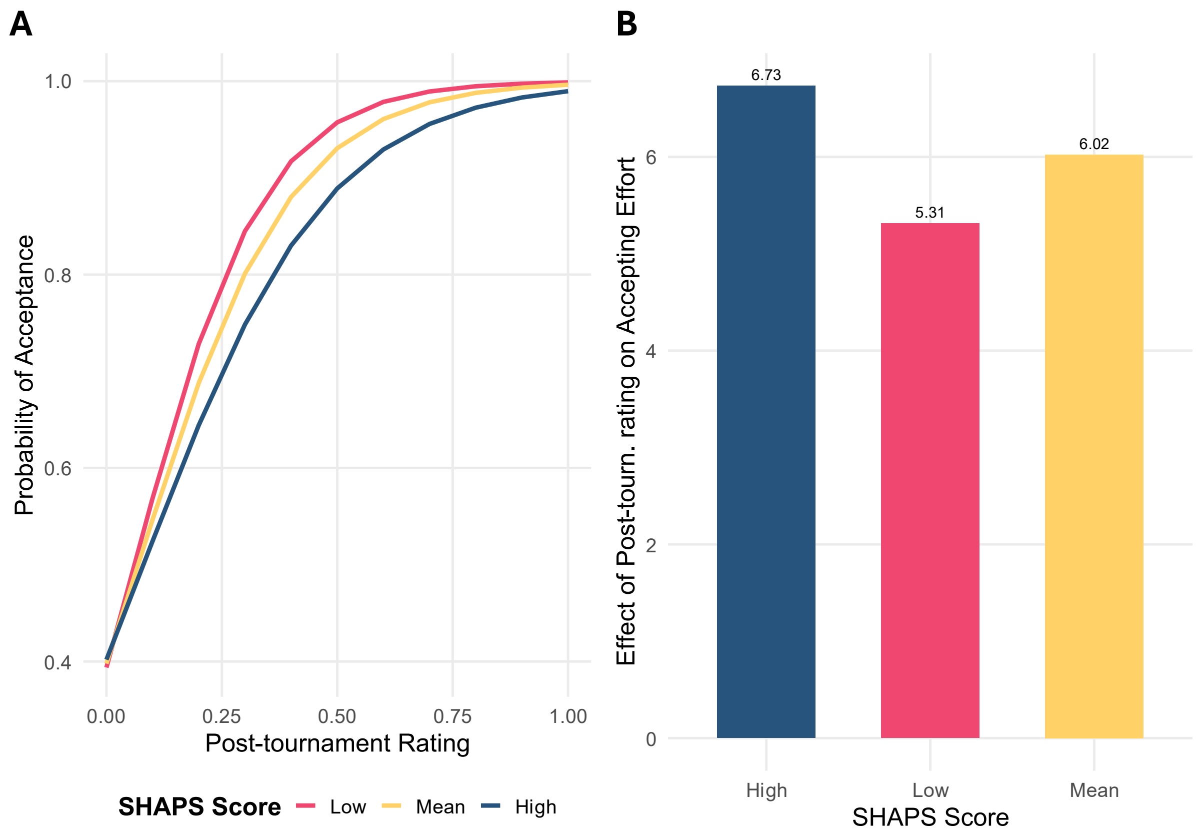


**Figure 6: The interaction between SHAPS scores and participants' post-tournament probability estimates. (A)** Predictions from our winning interaction model (EM1) that visualise the effect of the interaction between participants' post-tournament probability estimates and SHAPS scores on the probability of accepting effort offers. Compared to participants scoring lower on the SHAPS (cream colour), high SHAPS scorers' (red colour) effort choices are more weakly related to their estimates of the stimuli's outcome probability. Lower scorers on the SHAPS show a sharper increase in probability of accepting as their estimate of the outcome probability increases than participants who score higher on the SHAPS. **(B)** A bar plot of the interaction effect estimates for high, low and mean scorers on the SHAPS. These estimates were calculated by adding and subtracting the estimate for the interaction effect from the fixed effect estimate for participants' post-tournament estimates from the winning interaction model (EM1).

### Supplementary Materials

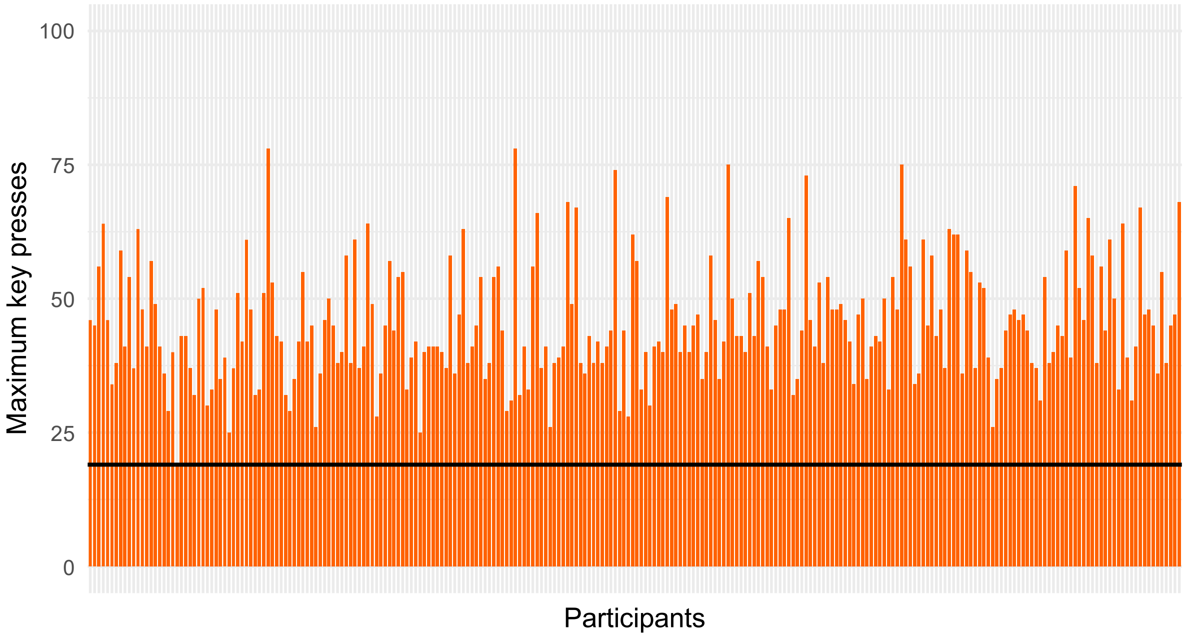


**Supplementary Figure 1: Bar plot of participants' maximum key presses from the effort calibration.** Each bar represents a participant's maximum number of key presses from the three calibration trials completed at the start of the experiment. The highest number of presses completed in the three calibration trials was taken forward to the effort decision-making task. Only one participant in the final sample had a maximum number of key presses equal to the minimum required to participate in the remainder of the experiment (19 presses).

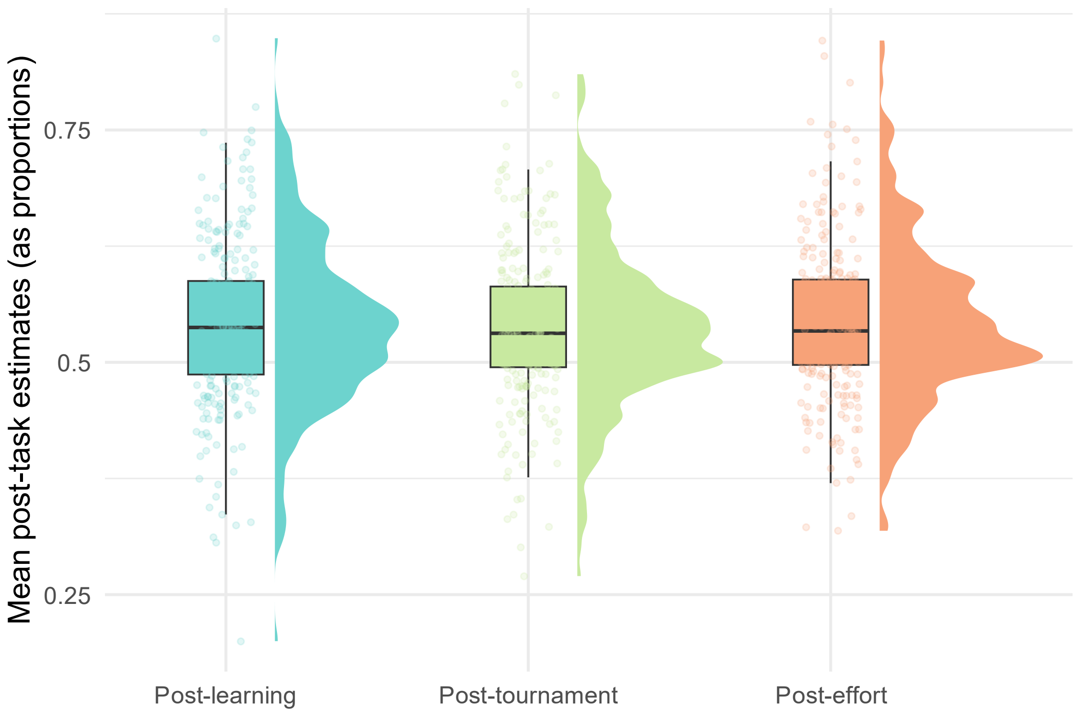


**Supplementary Figure 2: Mean post-task estimates across the experiment.** Rain cloud plots represent the distribution of participants' outcome probability estimates, which remain consistent across the experiment. The boxplot represents the 25th and 75th percentiles, and the middle line represents the median rating. Whiskers are 1.5 times the 25$^{th}$ and 75$^{th}$ percentiles.

***Supplementary Table 1***

**One sample t-tests of post-tournament probability estimates**

| Stimulus | Stimulus valence | Outcome probability | Mean of participant estimates | t-stat. (p-value Bonferroni corrected) | 95% CI |
|---|---|---|---|---|---|
| **Alien 1** | Reward | 0.25 | 0.296 | 3.37 (p = 0.014)* | 0.27 0.32 |
| **Alien 2** | Reward | 0.75 | 0.635 | -8.98 (p < 0.001)*** | 0.61 0.66 |
| **Alien 3** | Reward | 0.25 | 0.463 | 14.78 (p < 0.001)*** | 0.43 0.49 |
| **Alien 4** | Reward | 0.75 | 0.753 | 0.21 (p = 1) | 0.73 0.78 |
| **Alien 5** | Loss | 0.25 | 0.302 | 4.58 (p < 0.001)*** | 0.27 0.32 |
| **Alien 6** | Loss | 0.75 | 0.697 | -3.3 (p = 0.018)* | 0.67 0.73 |
| **Alien 7** | Loss | 0.25 | 0.417 | 11.61 (p = 0.001)** | 0.42 0.39 |
| **Alien 8** | Loss | 0.75 | 0.74 | -0.65 | 0.71 0.77 |

* p < 0.05 ** p < 0.01 *** p < 0.001

**Table S1: Summary of Bonferroni corrected one-sample t-tests comparing participants' post-tournament probability estimates against each of the stimuli's true probability.** The average of participants' post-tournament probability estimates for each stimulus demonstrates that participants tended to approximate the true outcome probabilities. Participants were most accurate in estimating the two stimuli with the highest absolute expected values (Alien 4 and Alien 8). Note that Bonferroni correction is applied by multiplying the p-value of each comparison by the number of comparisons, these adjusted p-values are then compared to the original significance threshold of 0.05. Asterisks therefore indicate whether a one sample t-test was significant at this threshold.

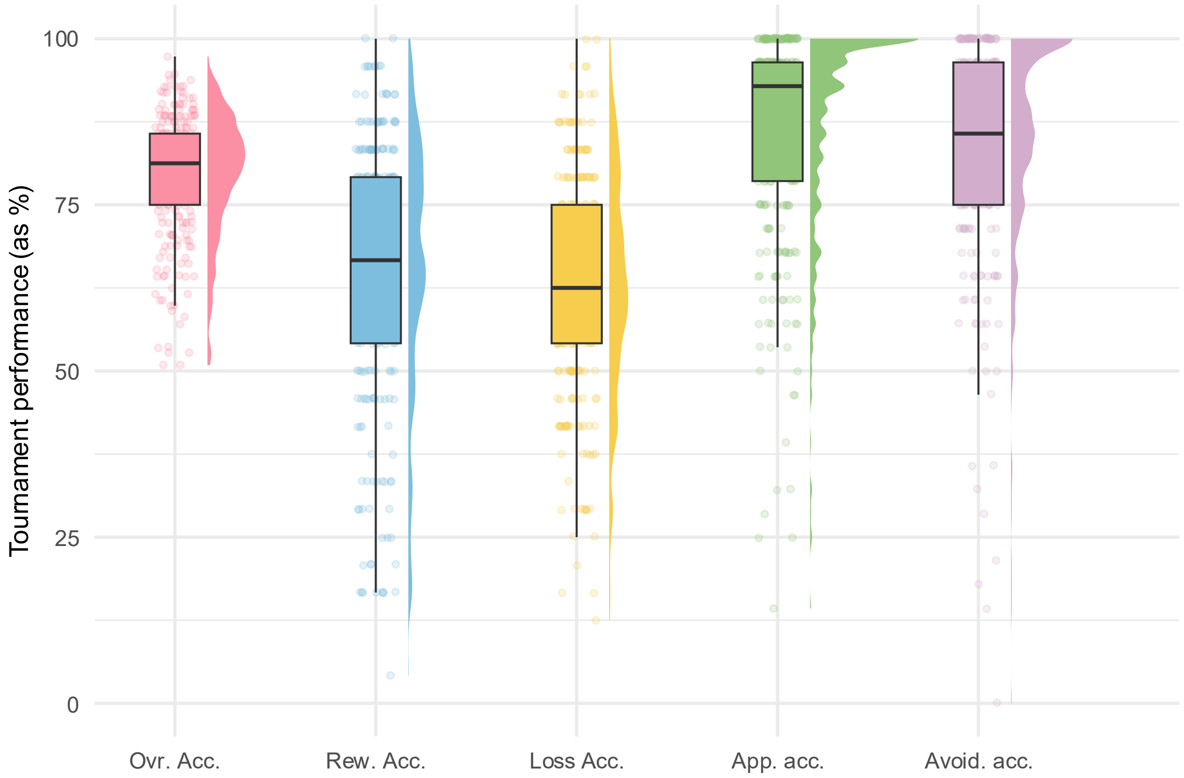


**Supplementary Figure 3: Rain cloud plots visualising participants' performance in the tournament task. From left to right: Overall accuracy** the percentage of trials in which participant chose the correct option, where the correct option was defined as the option which had the highest expected value (outcome magnitude * outcome probability); **Reward accuracy** the percentage of trials where participants chose the correct option when both options were stimuli associated with reward; **Loss accuracy** the percentage of trials where participants chose the correct option when both options were stimuli associated with loss; **Approach accuracy** the percentage of trials where participants chose the best stimulus (i.e. the stimulus with the overall highest expected value in the stimulus set) when it was present in a choice pair; **Avoid accuracy** the percentage of trials where participants chose any stimulus other than the worst stimulus (i.e. the stimulus with the lowest expected value in the stimulus set).

***Supplementary Table 2***

**Model 0 Summary**

| | Coefficient | Wald stat. (p value) | Tolerance | VIF |
|---|---|---|---|---|
| **Intercept** | 4.973 | 35.121 (p < 0.001) | | |
| **Effort level** | -7.004 | 46.966 (p < 0.001) | 0.98 | 1.031 |
| **Outcome probability** | 2.58 | 37.037 (p < 0.001) | 0.99 | 1.028 |
| **Outcome magnitude** | 0.774 | 13.921 (p < 0.001) | 0.99 | 1.006 |
| **Learning order (loss aliens first)** | -0.102 | 3.074 (p = 0.002) | 0.99 | 1.009 |
| **Valence (effect of reward)** | 0.224 | 7.743 (p < 0.001) | 0.99 | 1.001 |
| **Age** | 0.004 | 2.371 (p = 0.018) | 0.98 | 1.049 |
| **SES** | -0.008 | -0.866 (p = 0.386) | 0.98 | 1.049 |
| **Gender (male as reference):** | | | 0.99 | 1.022 |
| **Non-binary** | -1.06 | 19.844 (p < 0.001), | | |
| **Female** | 0.265 | 7.937 (p < 0.001), | | |
| **Not specified** | 0.78 | 2.518 (p < 0.011) | | |

**Table S2: Summary of the logistic regression model (M0) fitted to participants' trial-by-trial effort choices.** The overall model was significant ($X^2$ = 4248.173, $p < 0.001$) and effort level, outcome probability, outcome magnitude, learning order, valence and gender were significant predictors.

***Supplementary Table 3***

| Evaluation of mixed models containing individualised learning measures | | | | | | |
|---|---|---|---|---|---|---|
| **Method** | **Equation (in R syntax)** | **Number of parameters** | **AIC** | **BIC** | **Log likelihood** | **Deviance** |
| GLMM | choice ~ outcome prob. + outcome magn. + effort level + valence + age + gender + (1 + outcome prob. + outcome magn. + effort level + valence \| participant) | 20 | **14808** | 15002 | -7379.8 | 14760 |
| GLMM | choice ~ post-learning estimate + outcome magn. + effort level + valence + age + gender + (1 + post-learning estimate + outcome magn. + effort level + valence \| participant) | 20 | **14444** | 14638 | -7197.8 | 14396 |
| GLMM | **choice ~ post-tournament estimate + outcome magn. + effort level + valence + age + gender + (1 + post-tournament estimate + outcome magn. + effort level + valence \| participant)** | **20** | **13951** | **14145** | **-6951.4** | **13903** |
| GLMM | choice ~ mean trial-by-trial estimate + outcome magn. + effort level + valence + age + gender + (1 + mean prob. estimate + outcome magn. + effort level + valence \| participant) | 20 | **14722** | 14916 | -7337.1 | 14674 |
| GLMM | choice ~ final learning estimate + outcome magn. + effort level + valence + age + gender + (1 + mean prob. estimate + outcome magn. + effort level + valence \| participant) | 20 | **14756** | 14950 | -7354 | 14708 |

**Table S3 Comparison of the models which compare the predictive value of individual learning measures**. Table includes: model type, model equation in R (lme4) syntax, number of parameters, AIC, BIC, log likelihood and deviance scores for each model. These models were fit using *lme4* in R (R version 4.4.1; R Core Team, 2024; https://www.R-project.org/) package *lme4 (*version 1.1-35.5; Bates et al. 2024*)*. Details of the individualised learning measures can be found in the **Methods** section. Of these models, Model 3, which included participants' post-tournament probability estimates, was the best-fitting of all the individualised learning measures based on all model comparison metrics. Importantly, all models which contained an individualised learning metric outperformed the model (Model 1) which contained objective outcome probabilities.

##### ***Correlation amongst individualised learning measures***

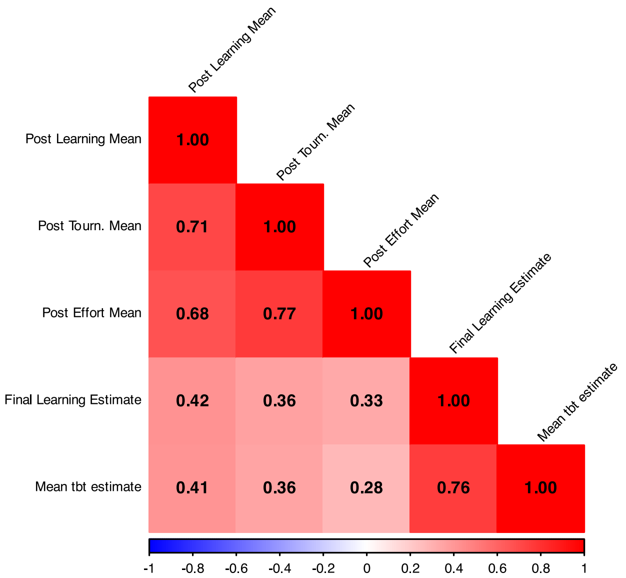


**Supplementary Figure 4: A correlation matrix of the individual learning measures.** Correlations amongst all five individualised learning measures used as substitutes for objective outcome probabilities in the generalised linear mixed effects models (GLMMs). All post-task probability estimates were at least moderately correlated with each other. These estimates were less strongly related to measures taken during the learning task (Final Learning Estimate and Mean tbt estimate), but these two measures were strongly associated with one another.

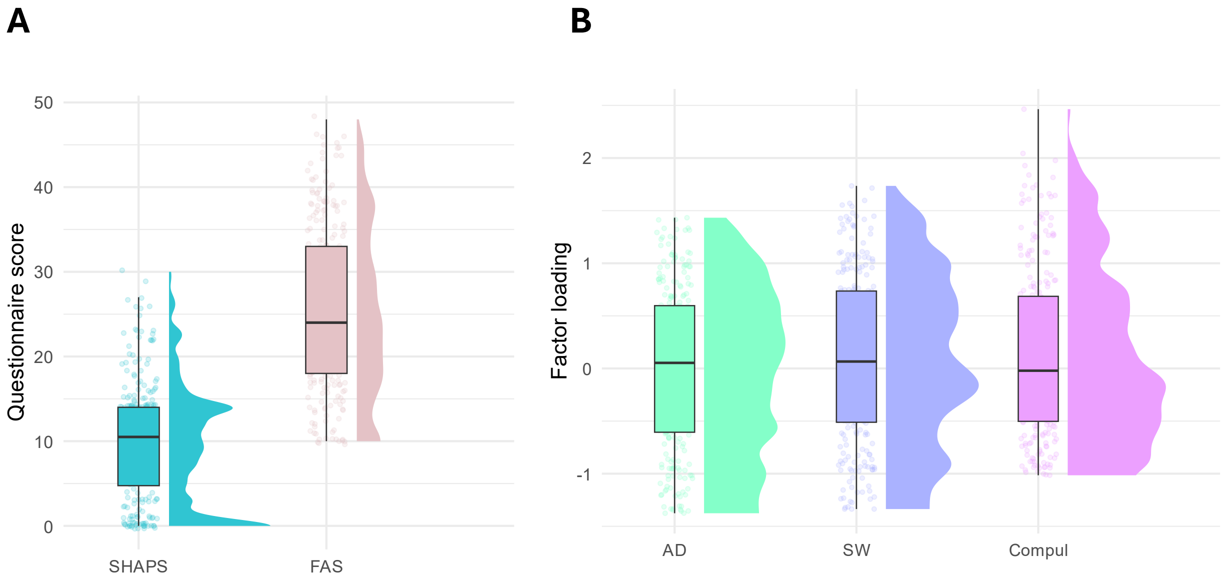


**Supplementary Figure 5: Anhedonia, fatigue scores and transdiagnostic factors. (A)** Raincloud plots of participants' summed scores on the SHAPS, a measure of anhedonia and the FAS, a measure of fatigue. **(B)** Raincloud plots of participants' loadings onto three transdiagnostic mental health factors, anxious-depression, social withdrawal and compulsivity (Gillan et al., 2016) using a reduced set of questionnaire items (Hopkins et al., 2022).

***Supplementary Table 4***

**Equations and evaluation of models with mental health predictors**

| Model number | Method | Equation (in R syntax) | Number of parameters | AIC | BIC | Log likelihood | Deviance |
|---|---|---|---|---|---|---|---|
| **0** | Logistic regression | choice ~ outcome prob. + outcome magn. + effort level + learning order + valence + gender + age + SES | 11 | **22468** | 22557 | -11223 | 22446 |
| **1** | GLMM | choice ~ post-tournament estimate + outcome magn. + effort level + valence + gender + age + (1 + post-tournament + outcome magn. + effort level + valence \| participant) | 24 | **13951** | **14145** | **-6951.4** | **13903** |
| **2** | GLMM | choice ~ post-tournament estimate + outcome magn. + effort level + valence + AD + Compul. + SW + gender + age + (1 + post-tournament estimate + outcome magn. + effort level + valence \| participant) | 27 | **13955** | 14173 | -6950.3 | 13901 |
| **3** | **GLMM** | **choice ~ post-tournament estimate + outcome magn. + effort level + valence + SHAPS + gender + age + (1 + post-tournament estimate + outcome magn. + effort level + valence \| participant)** | **25** | **13948** | **14150** | **-6948.9** | **13898** |
| **4** | GLMM | choice ~ post-tournament estimate + outcome magn. + effort level + valence + FAS + gender + age + (1 + post-tournament estimate + outcome magn. + effort level + valence \| participant) | 25 | **13952** | 14155 | -6951.2 | 13902 |

**Table S4: A summary of the models fitted to predict participants trial-by-trial choice behaviour** Including: fitting method used, the model equation written in the syntax of *lme4* in R (R version 4.4.1; R Core Team, 2024; https://www.R-project.org/) package *lme4 (*version 1.1-35.5; Bates et al. 2024*)*, the total number of model parameters, model AIC and BIC scores, model log likelihood and model deviance scores. Here we compare the performance of the three models which include mental health predictors, against the performance of a logistic regression (M0) and the winning model that did not include a mental health predictor (M1). Over and above Model 0 and Model 1, Model 3, which included the SHAPS as a fixed effect predictor, best fit participants' trial-by-trial choice data on all model comparison measures except BIC, which punishes more severely for additional model complexity.

***Exploratory analysis of the interaction between anhedonia and decision variables***

As our primary analysis identified SHAPS scores as a significant predictor of trial-by-trial effort choice, we conducted further analyses to explore whether SHAPS scores significantly interacted with participants' probability estimates, stimulus outcome magnitude, effort level or valence. For each interaction, we fit a separate exploratory model (EM) based on the winning model (M3):

$$\boldsymbol{EM1}: choice \sim outcome\ magnitude + effort\ level + stimulus\ valence + age + gender + \boldsymbol{SHAPS * individual\ learning\ measure} + (1 + individual\ learning\ measure + outcome\ magnitude + effort\ level + stimulus\ valence \mid participant)$$

$$\boldsymbol{EM2}: choice \sim individual\ learning\ measure + effort\ level + stimulus\ valence + age + gender + \boldsymbol{SHAPS * outcome\ magnitude} + (1 + individual\ learning\ measure + outcome\ magnitude + effort\ level + stimulus\ valence \mid participant)$$

$$\boldsymbol{EM3}: choice \sim individual\ learning\ measure + outcome\ magnitude + stimulus\ valence + age + gender + \boldsymbol{SHAPS * effort\ level} + (1 + individual\ learning\ measure + outcome\ magnitude + effort\ level + stimulus\ valence \mid participant)$$

$$\boldsymbol{EM4}: choice \sim individual\ learning\ measure + outcome\ magnitude + effort\ level + age + gender + \boldsymbol{SHAPS * stimulus\ valence} + (1 + individual\ learning\ measure + outcome\ magnitude + effort\ level + stimulus\ valence \mid participant)$$

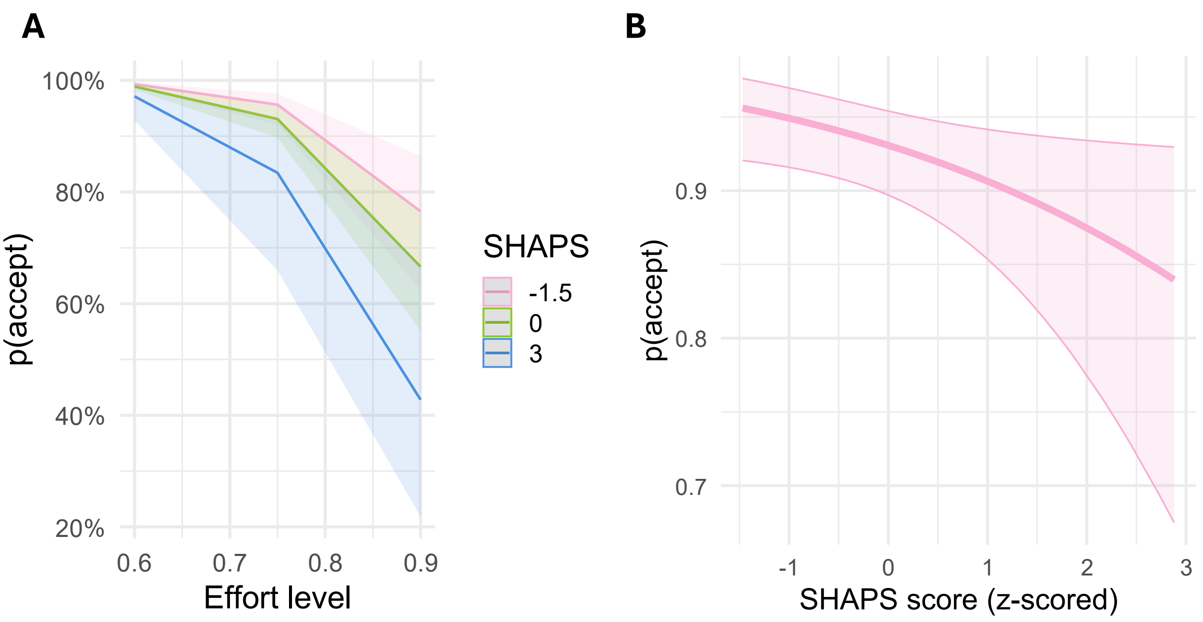


**Supplementary Figure 6: Line plots of the probability of accepting and SHAPS scores. (A)** A line plot representing predictions from the winning model (Model 3) of the effect of SHAPS scores (low, average and high scorers on the SHAPS) on the probability of accepting across effort levels. Please note this is only for visualisation purposes. Shaded areas are 95% CIs. **(B)** A line plot representing the fixed effect of SHAPS scores on the probability of accepting an effort offer. Shaded areas are 95% CIs.

***Supplementary Table 5***

| | | | | |
|---|---|---|---|---|
| **Reward stimuli** | | | | |
| **Alien** | 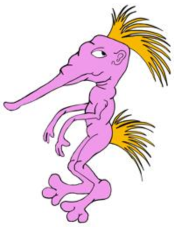 | 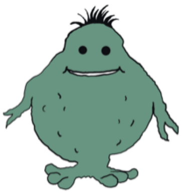 | 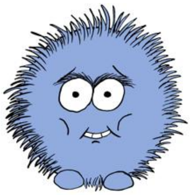 | 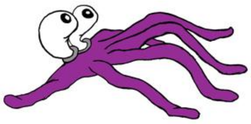 |
| **Outcome probability** | 0.25 | 0.75 | 0.25 | 0.75 |
| **Outcome magnitude** | 40 | 40 | 160 | 160 |
| **Loss stimuli** | | | | |
| **Alien** | 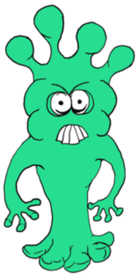 | 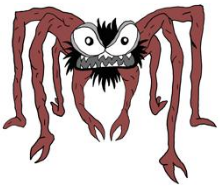 | 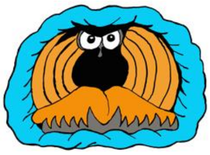 | 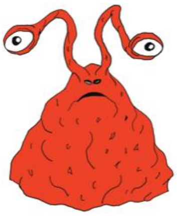 |
| **Outcome probability** | 0.25 | 0.75 | 0.25 | 0.75 |
| **Outcome magnitude** | -40 | -40 | -160 | -160 |

**Table S5: The eight aliens used for the learning, tournament and effort decision-making task.** Each alien is presented along with their respective outcome probabilities and outcome magnitude. Each stimulus was associated with a unique combination of outcome magnitude and outcome probability.
